# Molecular insights into the regulation of coenzyme selection in NADP-specific glutamate dehydrogenase from *Saccharomyces cerevisiae*

**DOI:** 10.64898/2026.11.02.763619

**Authors:** Arpan Banerjee, Alankar Roy, Spandan Saha, Barsa Kanchan Jyotshna Godsora, Prasenjit Bhaumik

## Abstract

Coenzyme specificity is one of the crucial aspects of the functional regulations in enzymes. Understanding the molecular basis of coenzyme specificity in glutamate dehydrogenase (GDH) is highly essential as this is the only enzyme that fulfils various physiological demands in terms of energy production and/or nitrogen assimilation via its strict functional regulation by connecting carbon and nitrogen metabolic cycles in every living organism. We have studied *S. cerevisiae* GDH3 (ScGDH3), as a model system for NADP-specific fungal GDHs. K272R and S250E mutants of ScGDH3 exhibited remarkable alteration in coenzyme specificity, with 15-55-fold and 3-14-fold more preference towards non-specific coenzymes [NAD(H)], respectively. S250E exhibited higher NADH affinity (∼ 1.3-fold) than that of towards NADPH. K272R showed ∼ 10-fold increased coenzyme affinity (*1/K_M_*) and binding affinity (*1/k_d_*) towards NADH, compared to the wild type enzyme (ScGDH3-WT). We solved the crystal structure of ScGDH3-K272R complexed with NADP^+^ at 2.50 Å resolution with a unique oligomeric arrangement having two distinct subunit conformations (“open” and “super-closed”) and it is the first structure of a GDH from *S. cerevisiae*. Molecular dynamics (MD) simulations provide insights into NADH binding and its increased affinity to ScGDH3-K272R. Further structure-based sequence analyses unravelled that a lysine and an arginine are naturally preferred residues for providing NADP(H) and NAD(H) specificity, respectively for various amino acid dehydrogenases. Taken together biochemical and structural data, our study successfully illustrates the evolutionary conserved mechanism of coenzyme related regulation in fungal and other GDHs; as well as amino acid dehydrogenases.

## Introduction

Enzymes are biological catalysts and they play crucial roles in various biochemical, physiological and metabolic processes of every living organism. Hence, the functions of enzymes are tightly regulated to maintain cellular as well as physiological homeostasis. Glutamate dehydrogenase (GDH) is one of the important metabolic enzymes present in all the living organisms. It is located in the cytosol of prokaryotic cells and in case of eukaryotic cells, GDH is generally present in mitochondria but the presence of GDH in the nucleus, lysosome and endoplasmic reticulum has also been reported (1). This oxidoreductase enzyme catalyses a reversible reaction that involves reductive amination of α-ketoglutarate (AKG) to form L-glutamate in presence of coenzyme NAD(P)H and ammonia and oxidative deamination of L-glutamate to form AKG in presence of coenzyme NAD(P)^+^ (Figure 1A). This reaction has tremendous physiological and metabolic importance as it interconnects the crucial carbon and nitrogen metabolic cycles (2, 3, 4). Formation of L-glutamate is an important step in nitrogen assimilation via several downstream processes and AKG being utilized via tricarboxylic acid (TCA) cycle aids in energy production for smooth functioning of cell. Malfunctioning of GDHs are often associated with several disorders such as hyperinsulinemia, propionic acidameia and tumour growth in human (5). In vertebrates, L-glutamate acts as neurotransmitter and cytoskeleton formation (6). It was reported that in plants and microbes, GDH is also involved in osmotic balance and high temperature tolerance (7). Hence, GDH plays a critical regulatory role to ensure balanced utilization of both L-glutamate and AKG based on the physiological need, to maintain the continuity of these two metabolic cycles at any cost.

**Figure 1:**
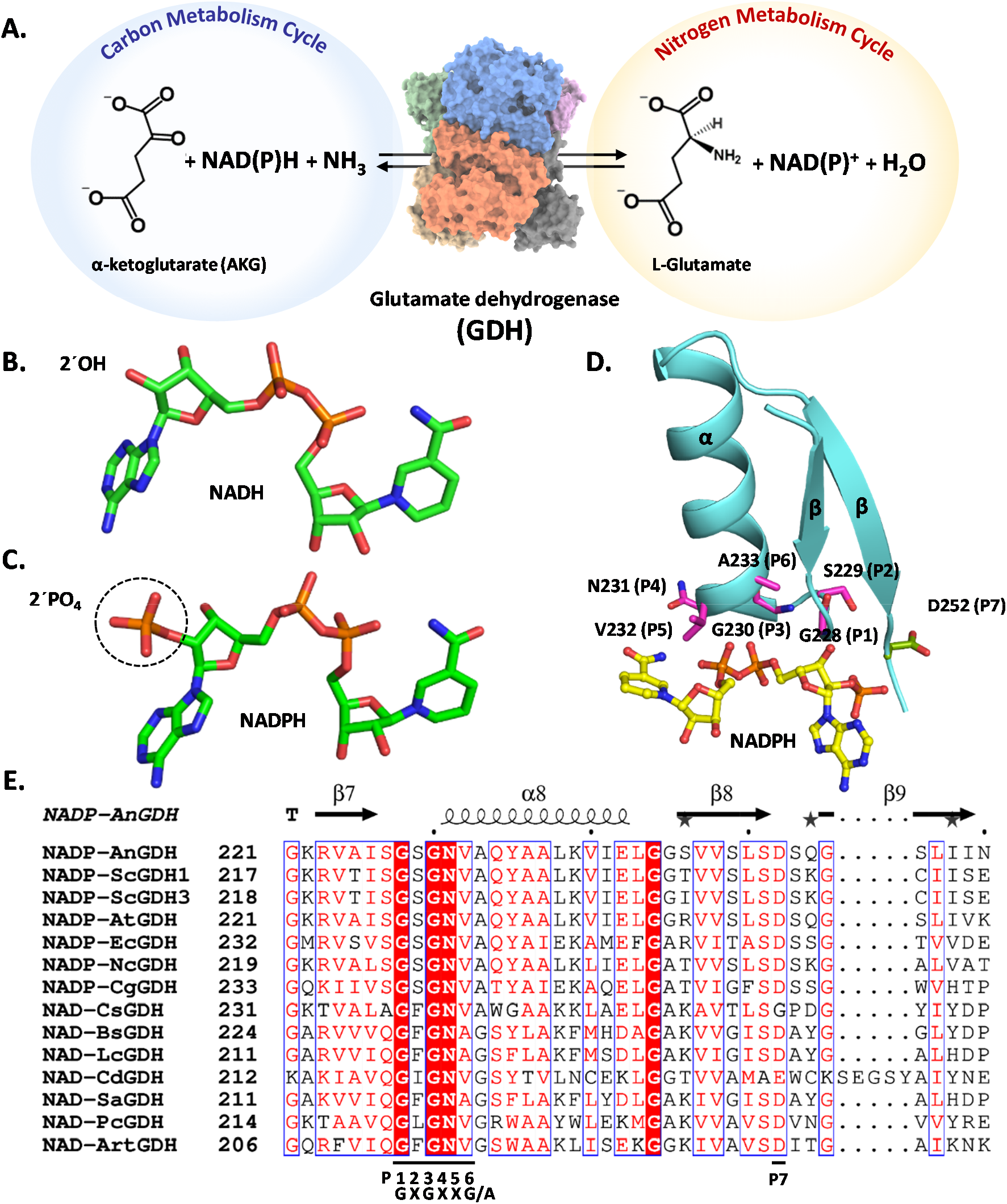
Catalytic reaction, coenzyme structures and coenzyme binding fold of GDHs. **A.** Reversible reaction including reductive amination (forward reaction) of AKG to form L-glutamate in presence of NAD(P)H as coenzyme and ammonia, as well as the oxidative deamination (reverse reaction) of L-glutamate to form AKG in presence of NAD(P) as coenzyme. AKG takes part in energy production via carbon cycle and L-glutamate helps in nitrogen assimilation. **B.** Structure of NADH with label on -2’OH associated with adenine ribose. **C.** Structure of NADPH with 2’PO4 moiety associated with adenine ribose highlighted by black dotted circle. **D.** Highly conserved β-α-β motif (represented in cartoon, aquamarine), part of the modified Rossmann fold, helps in coenzyme binding. The consensus “GXGXXG/A” sequence where residues (P1-P6) are shown in stick representation (magenta) which was proposed to stabilize coenzyme (Kleiger & Eisenberg, 2002); extremely conserved acidic residue (P7) (D252, shown in stick, smudge) is pointed away from NADPH. **E.** Selected portion of the sequence alignment using both NADP-specific and NAD-specific GDHs showing presence of the consensus “GXGXXG/A” sequence (P1-P6) and another extremely conserved acidic residue, named P7 (Baker et al., 1992). The sequence alignment was performed using GDHs from *Aspergillus niger* (AnGDH), *Saccharomyces cerevisiae* (ScGDH1 and ScGDH3), *Aspergillus terreus* (AtGDH), *Escherichia coli* (EcGDH), *Neurospora crassa* (NcGDH), *Corynebacterium glutamicum* (CgGDH), *Clostridium symbiosum* (CsGDH), *Bacillus subtilis* (BsGDH), *Lysinibacillus composti* (LcGDH), *Clostridioides difficile* (CdGDH), *Staphylococcus aureus* (SaGDH), *Pyrobaculum calidifontis* (PcGDH), and *Arabidopsis thaliana* (ArtGDH) respectively.

One of the important aspects of this layered functional regulation is coenzyme selection or coenzyme specificity. GDH mandatorily needs coenzymes for its catalytic reaction and two major coenzymes involved are nicotinamide adenine dinucleotide phosphate [NADP(H)] and/or nicotinamide adenine dinucleotide [NAD(H)] (Figure 1, B and C). Both the coenzymes are used either in their oxidised or reduced states depending on the nature of the reaction. In spite of having remarkable similarity in the structures with almost identical oxidation-reduction potential (8), they have quite different metabolic roles. In all the living organisms, NAD(H) is used as a catabolic cofactor whereas, NADP(H) has a predominant role in anabolic reactions such as biosynthesis (9). The same trend applies for GDH as well; NAD-specific GDHs are responsible for glutamate catabolism and NADP-specific GDHs catalyse glutamate biosynthesis via the utilization of AKG (10). Understanding the basis of coenzyme selection became essential to unravel the actual physiological role(s) of these enzymes. One such study revealed the presence of a consensus sequence “GXGXXG/A” (denoted as P1-P6) which is located in a β-α-β fold, present in domain II and help in recognition of adenine dinucleotide region of NAD(H) and NADP(H) (Figure 1, D and E) (11). The amino acid present in the P6 position, was identified as critical determinant of coenzyme specificity in different GDHs. Glycine (G) in the P6 position indicated NAD-specificity and alanine (A) in the same position indicated NADP-specificity (10). But, this rule had several exceptions and one of which was well characterized NAD-specific *Clostridium symbiosum* GDH (CsGDH) with alanine residue in the P6 position. Even the alanine to glycine mutation did not changed the overall coenzyme specificity in CsGDH (12). Another extremely conserved acidic residue (D/E) lining the coenzyme binding pocket, was identified and its position was named P7 (Figure 1E). The side chain orientation of this residue was found to be changed depending on the coenzyme molecule present at the pocket and hence considered crucial for determining coenzyme specificity (13, 14, 15) The evolutionary basis of coenzyme specificity unravelled the fact that both shared ancestry and sequence similarity are inadequate to explain identical coenzyme specificity (16). Rather, the coenzyme specificity may have evolved independently multiple times, with different structural strategies. It is difficult to explain the molecular basis with one universal structural mechanism (17). Extensive work primarily on bacterial NAD-specific GDHs revealed the possible molecular basis of coenzyme specificity and those studies showed that domain II alone is sufficient for selective and productive binding of coenzymes and thus, confers coenzyme specificity (18). In another study, the same group focused on the residues located in the coenzyme binding pocket and lining the cleft between two domains of CsGDH which strongly prefers NAD(H) as coenzyme. Based on the sequence comparison and pocket size analysis, they generated a triple mutant (F238S/P262S/D263K) that showed complete reversal of coenzyme specificity from NAD(H) to NADP(H) (12, 19, 20). Oliveira and his group generated a CEC chimeric GDH using domain I of NAD-specific CsGDH and domain II of *E. coli* GDH (EcGDH), an NADP-specific enzyme. The chimeric GDH showed NADP preference, similar to EcGDH. This further confirmed the coenzyme specificity is encoded by domain II residues in bacterial GDHs (21). Specific studies on dual coenzyme specific mammalian GDHs revealed the versatile role of P7 acidic residue. When NADP is present, it forms a salt-bridge with nearby positively charged residue which interacts with 2ʹPO_4_ moiety of NADP(H). In NAD-complexed structure, the same residue interacts directly with 2ʹOH of adenine ribose (14). However, the molecular basis of coenzyme specificity is very less explored in fungal GDHs mainly due to lack of coenzyme-complexed structures. Two recently solved ternary complex structures of NADP-specific GHDs from *Aspergillus niger* (AnGDH) and *Candida albicans* (CaGDH) revealed the presence of the consensus motif (“GXGXXG/A”) which stabilizes the coenzyme via H-bonding. Both of these studies revealed that the 2ʹPO_4_ moiety directly interacts with one lysine residue belongs to domain I. The fact these recent studies indicate that all the determinants of coenzyme specificity of GDHs might not exclusively belong to domain II (22, 23, 24) and domain I residues are also involved for recognising NADP(H). One recent study on *Schizosaccharomyces pombe* GDH1 (SpGDH1) revealed the possibility of phosphorylation-mediated regulation of this enzyme. They mimicked phosphorylation by mutating four different serine residues (S252, S281, S285, S287) located near the coenzyme binding pocket, which exhibited decreased NADP-specific activity. Based on the observations, they proposed a phosphorylation-mediated regulation of coenzyme specificity in SpGDH1(25). Despite these findings, a comprehensive study focused on coenzyme specificity of fungal GDHs is still missing. This inspired us to proceed with searching for an ideal fungal system where we can investigate the structural basis of coenzyme preference and our findings can be extrapolated to any other fungal GDHs.

*Saccharomyces cerevisiae* is a universal model organism for lower eukaryotic species. This organism is unique in many aspects. *S. cerevisiae* is one of the rarest organisms that has two distinct NADP-specific *gdh* genes, express two NADP-specific isoenzymes (ScGDH1 and ScGDH3) (10). The same organism has another GDH (ScGDH2) which is NAD-specific. Former two isoenzymes belong to S-GDH family and are reported to form a hexameric assembly whereas, the later one belongs to the L-GDH family and forms a tetrameric assembly. Hence, *Saccharomyces cerevisiae* has three GDHs which are structurally distinct and have different coenzyme preference with different subcellular localizations. All these aspects are associated with proper GDH function at the physiological environment, makes the organism ideal for studying enzyme functional regulation.

In this study, we have selected ScGDH3 as a model fungal GDH to identify the molecular determinants of NADP-specificity. With a particular focus on the binding pocket of 2ʹPO_4_ moiety of NADPH, several highly conserved residues were identified, mutated and extensive biochemical/kinetic characterizations were performed. Among them K272R and S250E single mutants showed significant alterations in the biochemical properties indicating their individual role in determining coenzyme specificity. We have solved a very well resolved crystal structure of K272R complexed with NADP^+^ at 2.50 Å. Structural analysis and molecular dynamics (MD) simulation studies successfully explained the molecular basis of alteration in biochemical properties. Targeted mutation in K119 from domain I revealed its crucial role in coenzyme selection and proper enzyme catalysis. This is the first comprehensive study on a fungal GDH unravelling the role of active site microenvironment which dictates the selection of appropriate coenzyme, crucial for the enzyme function and regulation. Taken together the biochemical and structural data, our study presents a solid foundation in understanding the molecular basis of coenzyme specificity in fungal GDHs.

## Results

### Specificity of the wild type ScGDH3 (ScGDH3-WT) and mutants towards NAD(P)H and NAD(P)^+^

For the mutagenesis study in ScGDH3, residues interacting with the 2’PO_4_ moiety of NADPH were initially selected. A total of five single mutants (K272R, K272A, S250E, K119A and K119F) and a double mutant (K272R-S250E) were generated. The rationales behind these mutations are discussed in detail in the “experimental procedures” section.

The enzyme activity assays were performed for both forward and reverse reactions catalysed by ScGDH3 using the wild type ScGDH3 (ScGDH3-WT) and all the mutant enzymes (Table 1, Figure 2). Specific activities for ScGDH3-WT with coenzyme NADPH and NADH were measured to be 46.50 (± 0.74) U/mg and 0.11 (± 0.01) U/mg, respectively (Figure 2, A and B). Specific activities for ScGDH3-WT with NADP^+^ and NAD^+^ were 5.07 (± 0.08) U/mg and 0.20 (± 0.001) U/mg, respectively (Figure 2, C and D). Significantly low specific activity for ScGDH3-WT was observed with coenzyme NAD(H) compared to that of with NADP(H), suggesting the native enzyme has strict and very high coenzyme preference towards NADP(H).

**Figure 2:**
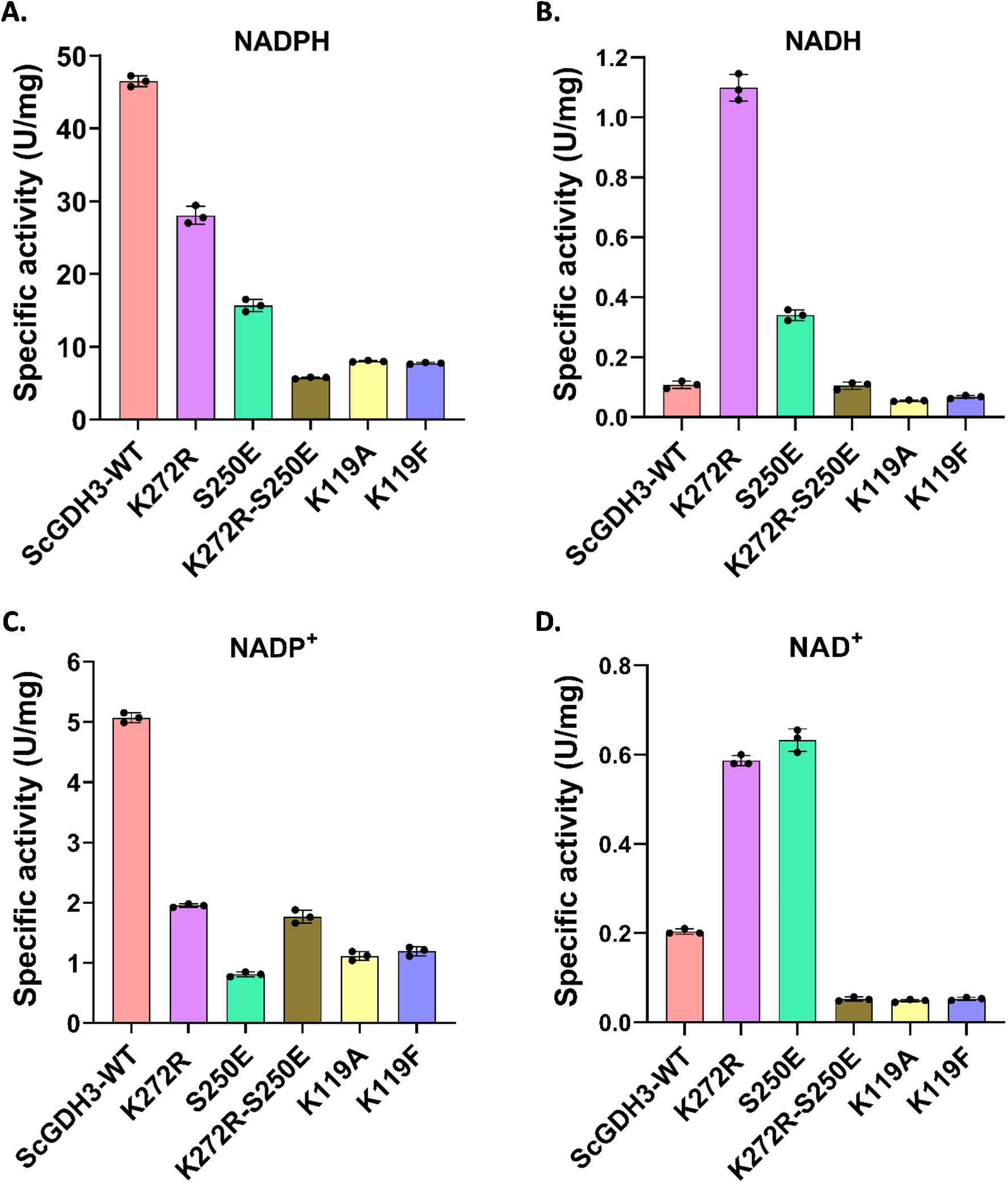
Bar graphs showing specific activities of ScGDH3-WT and the mutants. Specific activity of different enzyme variants when **A.** NADPH is considered as coenzyme, **B.** NADH is considered as coenzyme, **C.** NADP^+^ is considered as coenzyme, and **D.** NAD^+^ is considered as coenzyme. Specific activity of the mutants (K272R, S250E, K272R-S250E, K119A, K119F) are reported in μmoles/min/mg (U/mg). Among all the mutants, K272R and S250E exhibited significant amount of dual coenzyme specificity with lower NADP(H)-specific activity and higher NAD(H)-specific activity as compared to the native enzyme.

**Table 1:** Specific activities of ScGDH3-WT and all the mutants with NAD(P)H and NAD(P)^+^.

| **Enzyme** | **Specific activity (U/mg) with Coenzyme** |  |  |  |
|---|---|---|---|---|
|  | **NADPH** | **NADH** | **NADP<sup>+</sup>** | **NAD<sup>+</sup>** |
| **ScGDH3-WT** | 46.50 ± 0.74 | 0.11 ± 0.012 | 5.07 ± 0.08 | 0.20 ± 0.001 |
| **K272R** | 27.75 ± 1.68 | 1.08 ± 0.066 | 1.95 ± 0.03 | 0.58 ± 0.02 |
| **S250E** | 15.66 ± 0.84 | 0.34 ± 0.018 | 0.81 ± 0.04 | 0.63 ± 0.025 |
| **K272R-S250E** | 5.70 ± 0.12 | 0.11 ± 0.006 | 0.24 ± 0.04 | 0.052 ± 0.004 |
| **K119A** | 8.01 ± 0.18 | 0.056 ± 0.003 | 0.17 ± 0.02 | 0.049 ± 0.003 |
| **K119F** | 7.71 ± 0.13 | 0.068 ± 0.004 | 0.19 ± 0.02 | 0.053 ± 0.004 |

The K272R mutant exhibited dual coenzyme specificity to some extent (Figure 2, A-D). Specific activities for K272R mutant enzyme with coenzyme NADPH and NADH were measured 27.75 (± 1.68) U/mg and 1.08 (± 0.06) U/mg, respectively. K272R showed almost 1.68-fold reduction in NADPH-specific activity and 10-fold increase in NADH-specific activity compared to the native enzyme. Similar trend was observed when NADP^+^ and NAD^+^ were used and the specific activities were 1.95 (± 0.03) U/mg and 0.58 (± 0.02) U/mg, respectively. The K272R mutant introduced significant amount of NAD-specificity in a strictly NADP-specific enzyme (native). The K272A mutant showed negligible activity with both the coenzymes, indicating essentiality of K272 for the enzyme activity as well. All these observations proved the importance of K272 in regulating the coenzyme specificity in ScGDH3.

S250 residue was mutated to a negatively charged glutamate and activity of the S250E mutant was measured using both the coenzymes (Figure 2, A-D). The specific activities for S250E with NADPH and NADH were 15.66 (± 0.84) U/mg and 0.34 (± 0.02) U/mg, respectively. This mutant also exhibited dual coenzyme specificity with a significant reduction (∼ 3-fold) in NADPH-specific activity and increase (∼ 3-fold) in NADH-specific activity as compared to the native enzyme. Comparable specific activities were obtained when NADP^+^ and NAD^+^ were used as coenzymes. These data on the modulation of the activities in the S250E mutant clearly indicate the role of S250 in regulating coenzyme specificity in ScGDH3.

Based on the above observations, K272R and S250E mutations were combined and a double mutant (K272R-S250E) was generated. The specific activities for K272R-S250E with NADPH and NADH were 5.70 (± 0.12) U/mg and 0.11 (± 0.006) U/mg, respectively. The NADH-specific activity of this mutant is almost negligible and comparable to that observed for the native enzyme. In addition to that, NADPH-specific activity also reduced drastically (∼ 8-fold). Specific activities of the double mutant were also drastically reduced when NADP^+^ and NAD^+^ were used. It is evident that the double mutant becomes catalytically very less competent as compared to the native enzyme and the single mutants discussed so far.

The specific activities for K119F and K119A mutants with NADPH were 7.71 (± 0.13) U/mg and 8.01 (± 0.18) U/mg, respectively. Specific activities with NADH were 0.068 (± 0.004) U/mg and 0.056 (± 0.003) U/mg, respectively. For both these mutants, NADPH-specific activity reduced significantly along with almost zero NADH-specific activity. Exactly same trend was observed towards the NADP^+^ and NAD^+^ related specific activities for both of these mutants. Tryptophan fluorescence quenching assay was performed to check the effect of K119 residue in coenzyme binding. Interestingly, we observed almost similar extent of fluorescence quenching when NADPH (10-250 μM) binds with ScGDH3-WT and K119A mutant (Figure S1). Hence, K119A mutant severely affected enzyme activity but remain unaltered in terms of coenzyme binding. These observations revealed the importance of K119 in enzyme catalysis of ScGDH3.

### Enzyme kinetic assays of ScGDH3-WT and mutants with NAD(P)H and NAD(P)^+^

Coenzyme saturation kinetics were performed for both the forward and reverse reactions of ScGDH3-WT and two single mutants (K272R and S250E) with significant dual coenzyme specificity, using NAD(P)H and NAD(P)^+^. All the kinetic plots showed coenzyme saturation except for ScGDH3-WT with NAD(H) (Figures 3 and 4). This was expected as the native enzyme not at all prefers NAD(H) as coenzyme. All the kinetic parameters including enzyme turnover number (*k_cat_*) and catalytic efficiency (*k_cat_/K_M_*) are reported in Table 2 and 3.

**Figure 3:**
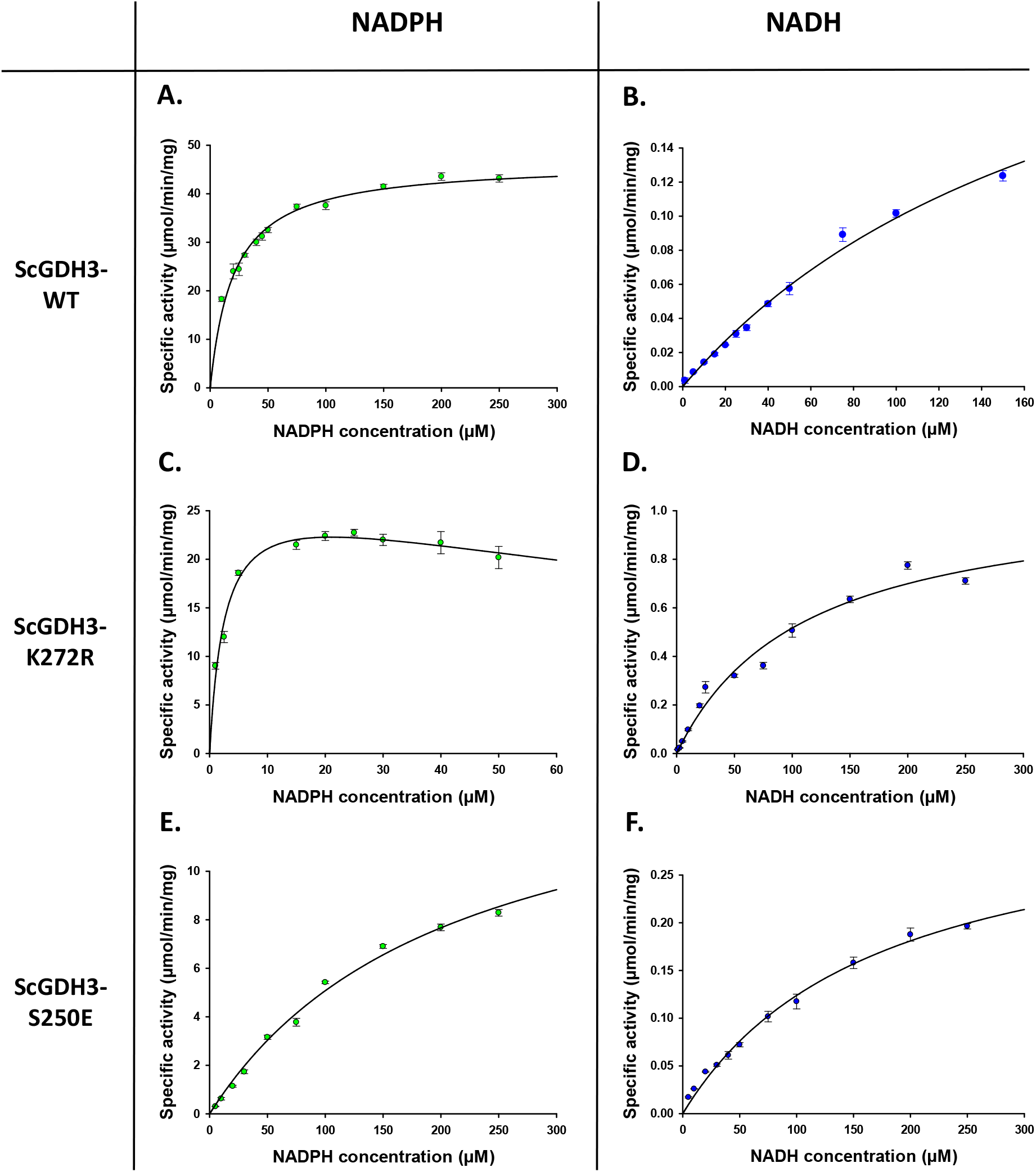
Coenzyme (NADPH and NADH) saturation kinetic plots of ScGDH3-WT and the mutants. **A.** NADPH saturation kinetics of ScGDH3-WT. **B.** NADH saturation kinetics of ScGDH3-WT. **C.** NADPH saturation kinetics of K272R mutant. **D.** NADH saturation kinetics of K272R mutant. **E.** NADPH saturation kinetics of S250E mutant. **F.** NADH saturation kinetics of S250E mutant. Specific activities are plotted against coenzyme [NAD(P)H] concentration to show saturation. ScGDH3 variants show different coenzyme saturation.

**Figure 4:**
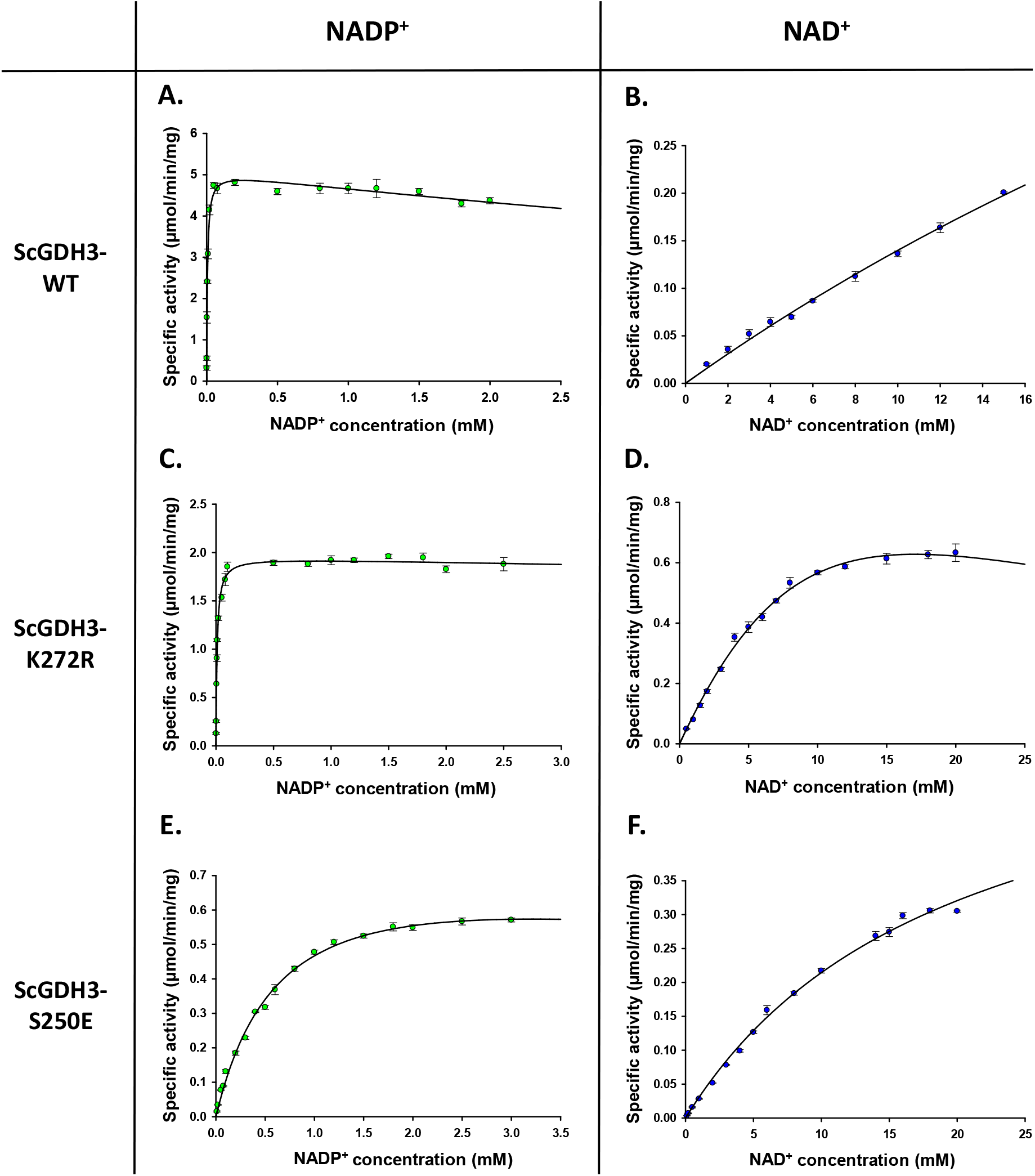
Coenzyme (NADP^+^ and NAD^+^) saturation kinetic plots of ScGDH3-WT and the mutants. **A.** NADP^+^ saturation kinetics of ScGDH3-WT. **B.** NAD^+^ saturation kinetics of ScGDH3-WT. **C.** NADP^+^ saturation kinetics of K272R mutant. **D.** NAD^+^ saturation kinetics of K272R mutant. **E.** NADP^+^ saturation kinetics of S250E mutant. **F.** NADP^+^ saturation kinetics of S250E mutant. Specific activities are plotted against coenzyme [NAD(P)^+^] concentration to show saturation. ScGDH3 variants show different coenzyme saturation.

**Table 2:** NAD(P)H saturation kinetic parameters of ScGDH3-WT, K272R and S250E mutants.

| **Enzyme** | **NADPH** |  |  |  | **NADH** |  |  |  | **Coenzyme preference<sup>#</sup>** |
|---|---|---|---|---|---|---|---|---|---|
|  | **V<sub>max</sub><br>(µmoles/min/mg)** | ***K<sub>M</sub>*** **(mM)** | ***k<sub>cat</sub>*** **(s<sup>-1</sup>)** | ***k<sub>cat</sub>/K<sub>M</sub>***<br>**(mM<sup>-1</sup>s<sup>-1</sup>)** | **V<sub>max</sub><br>(µmoles/min/mg)** | ***K<sub>M</sub>*** **(mM)** | ***k<sub>cat</sub>*** **(s<sup>-1</sup>)** | ***k<sub>cat</sub>/K<sub>M</sub>***<br>**(mM<sup>-1</sup>s<sup>-1</sup>)** |  |
| **ScGDH3-WT** | 46.50 ± 0.74 | 0.020 ± 0.001 | 38.33 ± 0.61 | 1916.5 ± 100.5 | 0.14 ± 0.025 | 0.204 ± 0.024 | 0.115 ± 0.02 | 0.56 ± 0.12 | 3422 |
| **K272R** | 27.75 ± 1.68 | 0.002 ± 0.0004 | 23.21 ± 1.40 | 11605 ± 2424.26 | 1.08 ± 0.06 | 0.11 ± 0.015 | 0.90 ± 0.05 | 8.18 ± 1.20 | 1419 |
| **S250E** | 15.66 ± 0.84 | 0.21 ± 0.018 | 13.03 ± 0.70 | 62.05 ± 0.63 | 0.34 ± 0.02 | 0.17 ± 0.016 | 0.286 ± 0.017 | 1.68 ± 0.19 | 37 |

\# These values represent the ratio of catalytic efficiency in the presence of NADPH (*k<sub>cat</sub>/K<sub>M</sub><sup>NADPH</sup>*) and NADH (*k<sub>cat</sub>/K<sub>M</sub><sup>NADH</sup>*) respectively, as coenzymes for individual enzymes.

Our biochemical studies showed few interesting phenomena, especially for the mutant K272R. This mutant exhibited drastically higher affinity (0.002 ± 0.0004 mM) towards NADPH compared to the native enzyme (0.020 ± 0.001 mM). This mutant also gained NADH affinity (0.109 ± 0.015 mM) and NAD^+^ affinity (6.75 ± 0.54 mM), unlike the native enzyme which has extremely low affinity towards these coenzymes (Figure S2). Most importantly, the catalytic efficiency of the K272R mutant with the non-specific coenzymes (NADH and NAD^+^) has increased by ∼ 15-fold and 55-fold respectively, as compared to the native enzyme. The significant alteration in coenzyme binding and kinetics properties is quite unique for a single point mutant (K272R). Therefore, we determined the crystal structure (as discussed in the later sections) of this mutant to understand the molecular basis of coenzyme recognition.

Kinetic studies also revealed significant alteration in the biochemical properties in S250E mutant as compared to the native enzyme. Very high *K_M_* value (0.21 ± 0.018 mM) indicates significantly lower affinity of S250E mutant towards NADPH. This mutant also gained affinity towards NADH (0.17 ± 0.016 mM) like the previous single mutant (K272R). S250E mutant also exhibited significantly higher (∼ 4-fold) NAD^+^-affinity (Figure S2). The catalytic efficiency of S250E mutant with the non-specific coenzymes (NADH and NAD^+^) has significantly increased by 3-fold and ∼ 14-fold respectively when compared with the native enzyme. These findings clearly indicate the S250E mutation exhibits more rigorous dual coenzyme specificity, indicating the crucial role of S250 in selecting appropriate coenzyme for ScGDH3.

### Binding affinity towards NAD(P)H

Binding affinities of ScGDH3-WT and K272R mutant towards NADPH and NADH were determined using MicroScale Thermophoresis (MST) spectral shift analysis. The dose response curves with ratio of fluorescence intensities at 670 nm and 650 nm against the coenzyme concentration are presented in (Figure 5, A-D). All the binding assays were performed at the room temperature and *k_d_* values were are reported in Table 4. Notably, the data showed significantly low noise level (0.002-0.004) and very high signal to noise ratio (9-13), indicating the high quality as well as reliability of these experiments. For ScGDH3-WT, the *k_d_* value towards NADPH was determined to be 1.42 ± 0.20 μM which indicates significant high affinity of the enzyme towards NADPH, as it is the preferred coenzyme. However, no spectral shift was observed in the dose response curve when NADH was used. The analysis clearly indicated very poor or no binding of NADH to the native enzyme. This result further proved that ScGDH3-WT is strictly NADPH-specific, NADH can’t even bind to it properly. K272R showed drastically high binding affinity towards NADPH with a much lower *k_d_* value of 0.13 ± 0.018 μM. This affinity is ∼ 10-fold higher than that of observed for the native enzyme towards NADPH (Figure 5E). K272R mutant also has quite high binding affinity towards NADH as well (*k_d_* = 12.90 ± 2.45 μM), indicating much tighter NADH binding unlike the native enzyme where no measurable NADH binding was detected (Figure 5F).

**Figure 5:**
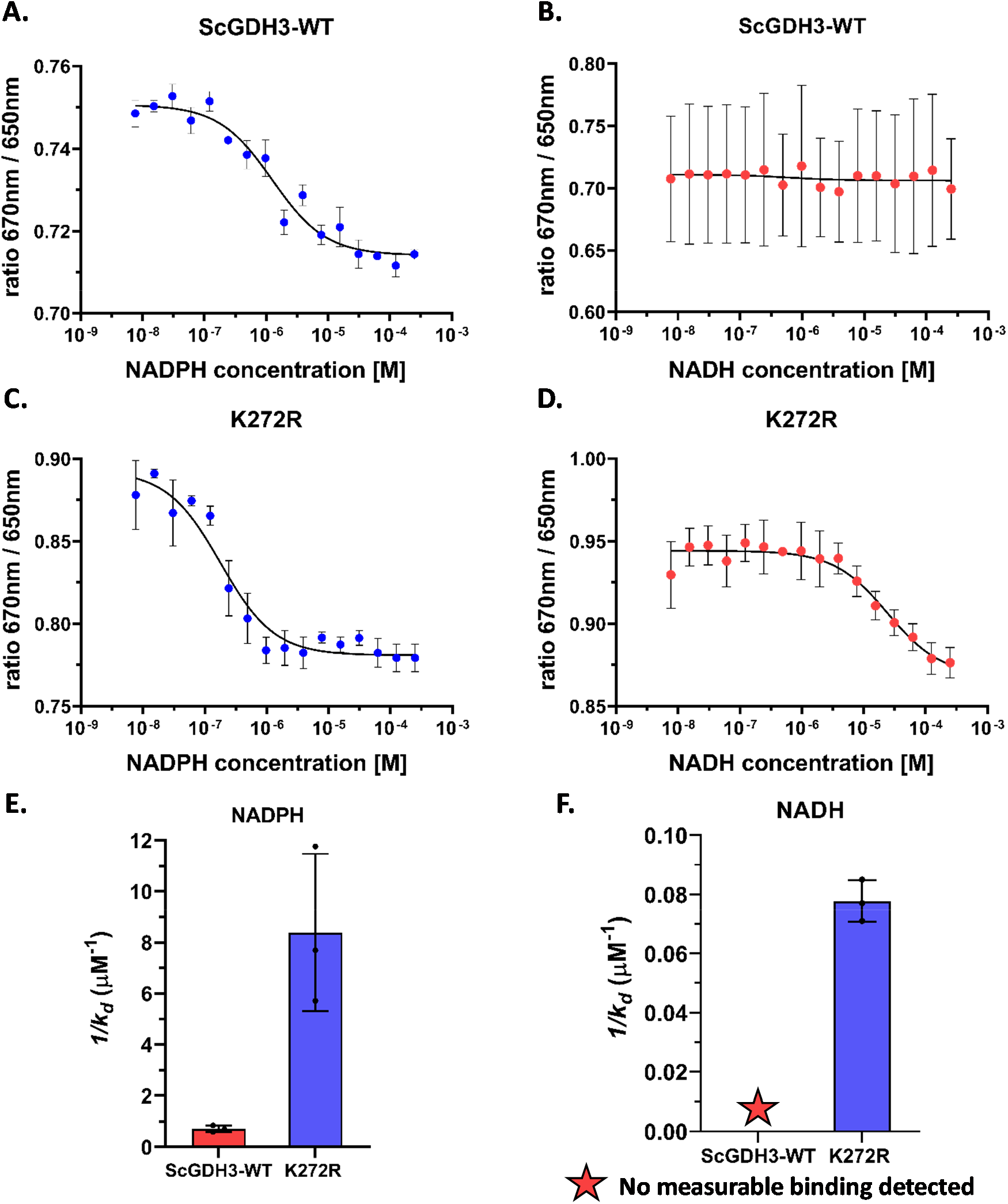
Spectral shift analysis showing coenzyme binding with ScGDH3-WT and K272R mutant. **A.** Binding of NADPH with ScGDH3-WT, **B.** Binding of NADH with ScGDH3-WT, **C.** Binding of NADPH with K272R mutant, and **D.** Binding of NADH with K272R mutant. The ratio of fluorescence counts at 670 nm and 650 nm was plotted against respective coenzyme concentration. Black line represents the dose response curve. The straight line indicates no binding of NADH with ScGDH3-WT which is strictly NADP(H)-specific. **E.** Bar graph showing binding affinity of ScGDH3-WT and K272R mutant towards NADPH. Y-axis presents reciprocal of dissociation constant (*1/k_d_*), suggesting greater the value higher will be the binding affinity towards a particular ligand. **F.** Bar graph showing binding affinity of ScGDH3-WT and K272R mutant towards NADH.

**Table 3:** NAD(P)^+^ saturation kinetic parameters of ScGDH3-WT, K272R and S250E mutants.

| **Enzyme** | **NADP<sup>+</sup>** |  |  |  | **NAD<sup>+</sup>** |  |  |  | **Coenzyme preference<sup>#</sup>** |
|---|---|---|---|---|---|---|---|---|---|
|  | **V<sub>max</sub> (µmoles/min/mg)** | ***K<sub>M</sub>* (mM)** | ***k<sub>cat</sub>* (s<sup>-1</sup>)** | ***k<sub>cat</sub>/K<sub>M</sub>* (mM<sup>-1</sup>s<sup>-1</sup>)** | **V<sub>max</sub> (µmoles/min/mg)** | ***K<sub>M</sub>* (mM)** | ***k<sub>cat</sub>* (s<sup>-1</sup>)** | ***k<sub>cat</sub>/K<sub>M</sub>* (mM<sup>-1</sup>s<sup>-1</sup>)** |  |
| **ScGDH3-WT** | 5.07 ± 0.08 | 0.006 ± 0.0004 | 4.24 ± 0.07 | 706.67 ± 48.50 | 0.20 ± 0.027 | 72.28 ± 19.86 | 0.17 ± 0.023 | 0.002 ± 0.0006 | 353335 |
| **K272R** | 1.95 ± 0.03 | 0.009 ± 0.0006 | 1.62 ± 0.02 | 180.55 ± 12.35 | 0.89 ± 0.03 | 6.75 ± 0.54 | 0.75 ± 0.025 | 0.11 ± 0.009 | 1641 |
| **S250E** | 0.81 ± 0.04 | 0.68 ± 0.06 | 0.67 ± 0.03 | 0.98 ± 0.09 | 0.63 ± 0.025 | 19.58 ± 1.30 | 0.525 ± 0.021 | 0.027 ± 0.002 | 36 |

\# These values represent the ratio of catalytic efficiency in the presence of NADP<sup>+</sup> (*k<sub>cat</sub>/K<sub>M</sub><sup>NADP+</sup>*) and NAD<sup>+</sup> (*k<sub>cat</sub>/K<sub>M</sub><sup>NAD+</sup>*) respectively, as coenzymes for individual enzymes.

**Table 4:** *k_d_* values of NAD(P)H for ScGDH3-WT and K272R mutant.

| **Enzyme** | ***k<sub>d</sub>*** **(μM)** |  |
|---|---|---|
|  | **NADPH** | **NADH** |
| **ScGDH3-WT** | 1.42 ± 0.24 | - |
| **K272R** | 0.13 ± 0.045 | 12.90 ± 1.08 |

### Overall structural architecture of ScGDH3-K272R

We determined the first experimental 3D structure of a *S. cerevisiae* GDH. The crystal structure of ScGDH3-K272R complexed with NADP^+^ was solved at 2.50 Å resolution to decipher the mechanism of coenzyme specificity in ScGDH3 (Figure 6). The structure is well refined with acceptable parameters reported in Table 5. The excellent quality of the reported structure can be assessed by the well-defined electron density and very good statistics of the data collection as well as refinement. The structure of ScGDH3-K272R has been determined with four subunits (two dimers) in the asymmetric unit (Figure S3A). But, it forms the crystallographic hexameric (biological) assembly (Figure S3, B and C; Media S1). The functional homo-hexameric state of this enzyme is similar to other reported GDH structures from *E. coli*, *C. symbiosum* and *A. niger*. The hexameric assembly has 2-fold and 3-fold symmetries; and the structure can be described either as dimer of trimers or as trimer of dimers (23). The hexameric enzyme has an overall cylindrical shape with estimated height and width of 110 Å and 91 Å respectively (Figure 6, A-C). Each subunit is composed of two domains: domain I and domain II. There is a cleft present in between two domains where the active site of the enzyme is present (Figure 6D). The overall structural fold is found to be conserved in each of the subunits. The first N-terminal residue (M1) of each subunit couldn’t be built due to missing electron density. Domain I consist of residues 1-187 and 433-457, and domain II consists of residues 188-432. So, both N-terminal and C-terminal are part of domain I. Both α-helices and β-sheets have comparable contributions in the architecture of these domains (Figure 6E). Domain I starts from the N-terminus and continues till 187th residue, then domain II continues and ends before the C-terminal helix (α_16_). This helix protrudes towards domain I and considered part of domain I. The well-established modified Rossmann fold (β-α-β fold) is present in domain II. In the hexameric structure, domain I is found to be interacting with other subunits and plays an important role in the formation of hexameric assembly whereas domain II is majorly responsible for coenzyme binding. The dimeric interface is mainly formed by residues present in domain I. The residues from both domain I (α_3_ and α_16_ helices) and domain II (α_14_, hinge helix) lined the trimeric interface. Hence, majorly the domain I residues aid in the formation of the inter-subunit interfaces and maintain the hexameric assembly of ScGDH3-K272R as observed for other hexameric GDHs (22, 26).

**Figure 6:**
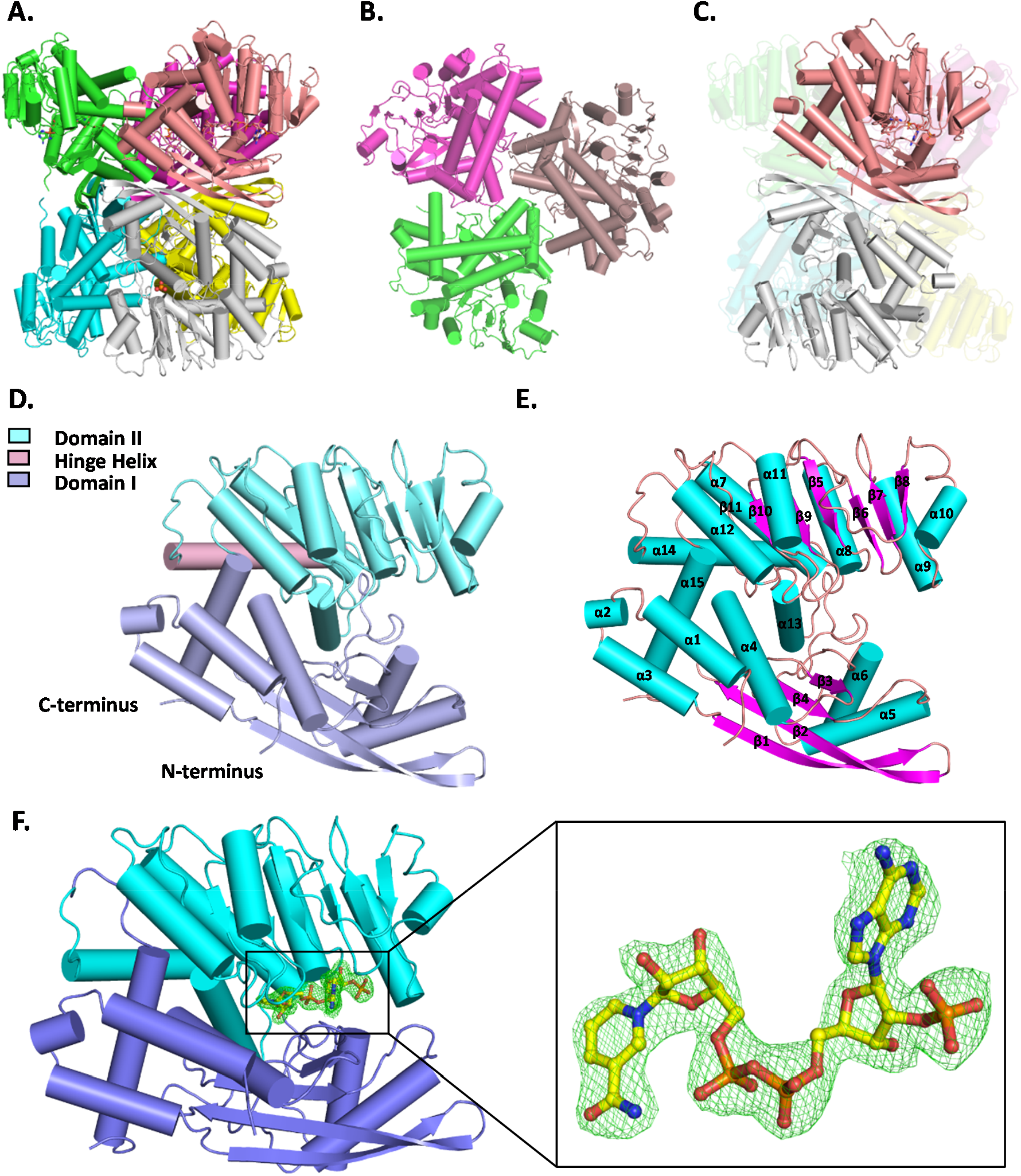
Overall structural fold of ScGDH3-K272R-NADP complex. **A.** Classic crystallographic hexameric assembly (functional state) represented in cartoon with cylindrical helices. Each subunit is colored differently. **B.** Top view of the hexameric assembly showing one trimeric (three subunits) interface region. **C.** One subunit from each trimer together forming a dimer; hexameric assembly can be represented as trimer of dimers or dimer of trimers. **D.** Tertiary structure of a single subunit composed of two different domains: domain I (blue white) and domain II (aquamarine). The hinge helix (dirty violet), part of domain II, has important role in regulating domain movements. **E.** The secondary structural elements present in every subunit are shown in cartoon: helices are labelled as “α” (cyan), sheets are labelled as “β” (magenta) and loops are colored as light pink. **F.** The coenzyme, NADP^+^ bound to the coenzyme binding pocket. Zoomed in view showing *F_o_-F_c_* omit map (represented as mesh in light green) for NADP^+^ contoured at 3.0 σ level.

**Table 5:** Data collection and refinement statistics of ScGDH3-K272R-NADP complex.

| **Data collection statistics<sup>#</sup>** | **ScGDH3-K272R-NADP** |
|---|---|
| Space group | *H* 3 2 |
| Unit cell parameters |  |
| *a, b, c* (Å) | 157.11, 157.11, 515.42 |
| *α,* β, γ (°) | 90.0, 90.0, 120.0 |
| Temperature (K) | 100 |
| Wavelength (Å) | 0.97893 |
| Resolution (Å) | 50.0-2.5 (2.6-2.5) |
| R<sub>merge</sub> (%) | 35.9 (293.3) |
| Completeness (%) | 99.6 (100.0) |
| Mean *I*/σ*I* | 7.65 (1.05) |
| Unique reflections | 84638 (9310) |
| Redundancy | 11.23 (11.31) |
| CC<sub>1/2</sub> (%) | 99.3 (49.8) |
| **Refinement statistics** |  |
| Resolution (Å) | 50.0-2.5 |
| No. of reflections (Working set/Test set) | 7966/418 |
| *R<sub>work</sub>* (%) | 20.8 |
| *R<sub>free</sub>* (%) | 26.5 |
| **Model composition** |  |
| Protein chains | 4 |
| Protein residues | 1824 |
| Number of NADP molecules | 2 |
| Number of water molecules | 337 |
| **Geometry statistics** |  |
| RSMD (bonds) (Å) | 0.008 |
| RMSD (angles) (°) | 0.952 |
| Ramachandran plot |  |
| Favoured (%) | 94.55 |
| Allowed (%) | 4.68 |
| Outliers (%) | 0.77 |
| **PDB ID** | 47UL |

\# values in parentheses correspond to the highest resolution shell

### Active site of ScGDH3-K272R and NADP^+^ binding

The crystal structure of NADP^+^-bound ScGDH3-K272R has four subunits (two dimers) in the asymmetric unit. One subunit of each dimer has two ligands bound to the active site. Interestingly, the crystallographic hexameric (biological) assembly has half (A, B and C; upper trimer) of the six subunits is complexed with the coenzyme NADP^+^ (Figure 6F) at the coenzyme binding pocket and a citrate (Figure S4A) at the substrate binding site. The well resolved electron density map clearly shows the presence of citrate (Figure S4A) and NADP^+^ (Figure 6F) at the active site of the subunits with “super-closed” conformation. The clearly visible positive *F_o_-F_c_* omit electron density (Figure S4B) indicates mutation of lysine to arginine at 272^nd^ position.

Two domains in each subunit oriented in such a way that it forms a cleft in between them. The active site is present at the core of this cleft. The citrate molecule (possibly from the mother liquor as discussed later) is found to be stabilized by the residues (K75, Q96, K99, K111, D151, R190 and S382) lining the substrate binding site (Figure S4C). Similar binding modes of other carboxylate group containing metabolites in the substrate (α-ketoglutarate) binding pocket have been reported for *Aspergillus terreus* GDH (27). Multiple residues are involved in the interaction with NADP^+^ to stabilize it in the specific site (Figure S5, A-E). Although most of them are from domain II, but domain I residues (R79, H81, K119, D151) also contributed significantly to accommodate NADP^+^ inside the binding pocket. Four residues (K119, D249, S250 and R272) were involved in the interaction (direct/indirect) with the 2’PO_4_ group of adenine ribose moiety of NADP^+^ (Figure 7A). All the residues are highly conserved among other NADP-specific GDHs, except the mutated R272 (lysine in the native enzyme is also conserved). K119 is the only residue from domain I that directly interacts with the 2’PO_4_ of NAPD^+^ (Figure 7B). S250 residue also involved in direct interaction with the 2’PO_4_ of NAPD^+^ (Figure 7C). The highly conserved D249 (the P7 residue) pointed away from the 2’PO_4_, which is evident in other NADPH-bound GDH structures (23, 24, 14). However, we also found a unique water mediated interaction between D249 and 2’PO_4_ (Figure 7D), not reported in any other previous structures. The arginine (R272) residue exhibited more focused and stringent interactions only with 2’PO_4_ (Figure 7E), unlike the lysine residue (in ScGDH3-WT). R272 residue also interacts with the 3’OH group of adenine ribose, which is unique and plays a critical role in increasing coenzyme affinity. Citrate in the substrate binding site, is majorly stabilized by domain I residues while NADP^+^ interacting residues are mostly from to domain II. However, residues like D151 and R190 were found to interact with both citrate and NADP^+^ in the active site. The subunit conformation (discussed in the next section) also aids in stabilizing NADP^+^ via the formation of appropriate coenzyme binding pocket.

**Figure 7:**
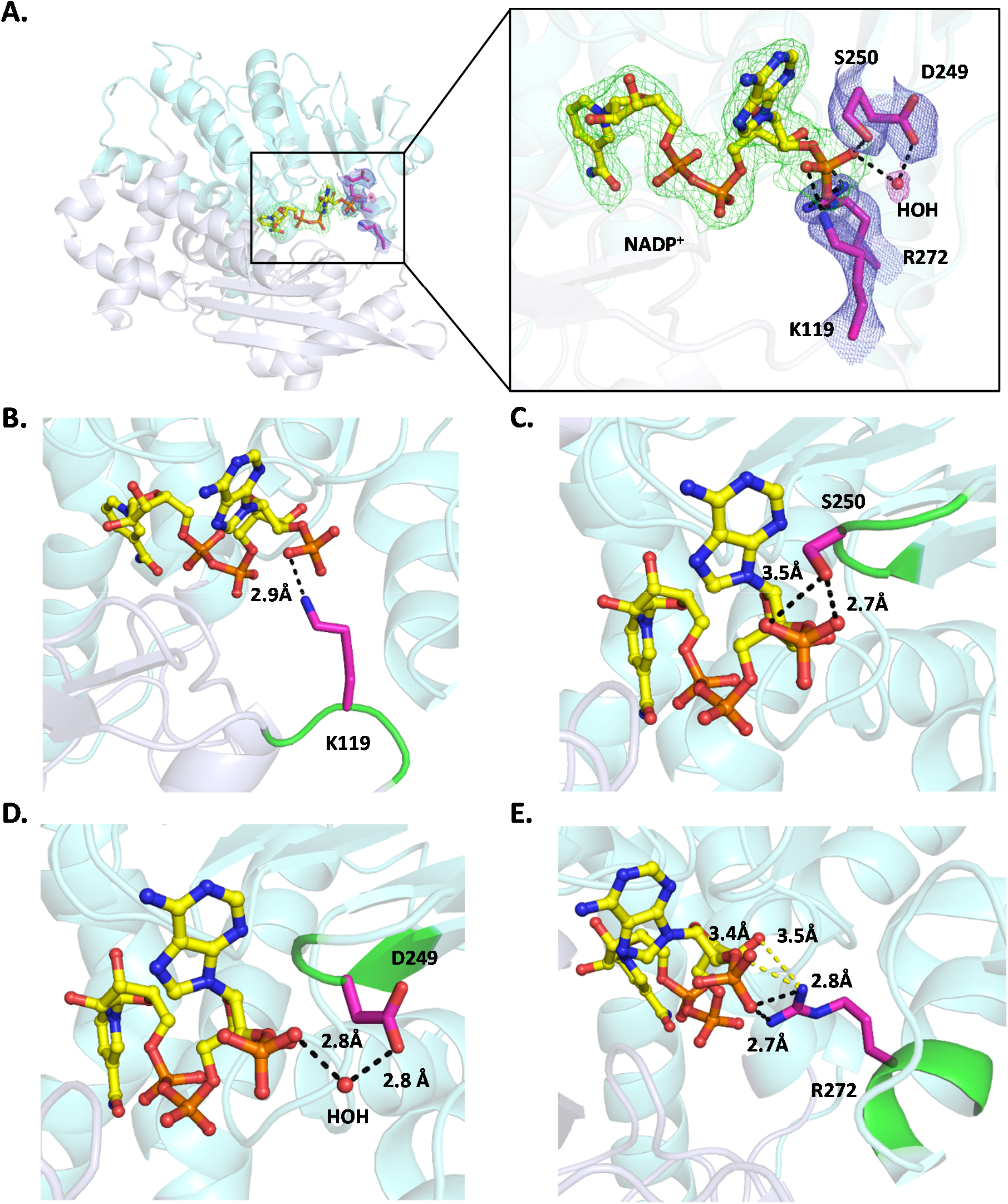
Molecular basis of binding of the 2’PO_4_ moiety of NADP^+^. **A.** Coenzyme bound subunit shown in cartoon consists of domain I (blue white) and domain II (aquamarine). Zoomed in view showing four residues (K119, D249, S250 and R272) represented in sticks (magenta), interact (direct/indirect) with 2’PO_4_ moiety of NADP^+^ (ball and sticks, yellow). The *2F_o_-F_c_* maps contoured at 1.0 σ level represented in mesh, showing NADP^+^ (green), interacting residues (slate) and water (light pink). **B.** K119 is directly involved in an electrostatic interaction with the 2’PO_4_ moiety. Green colour region of the cartoon represents the secondary structure in which the residues are located and black dotted line represents the interaction distance (labelled in Å). **C.** S250 also directly involved in polar interactions with 2’PO_4_ moiety. **D.** D249 is pointed away from 2’PO_4_ moiety and involved in a unique water (shown in sphere, red) mediated hydrogen bonding interaction with 2’PO_4_ moiety. **E.** The mutated residue R272 is involved in couple of polar interactions not only with 2’PO_4_ moiety but also with 3’OH group of adenine ribose as well. Yellow dotted lines represent the interaction distance (labelled in Å) other than the 2’PO_4_ moiety.

### Conformational heterogeneity of ScGDH3-K272R

GDHs are intrinsically very dynamic in nature (24, 26). The domain II is reported to have more flexibility as compared to domain I. The conformational dynamicity of domain II is associated with ligand binding and is regulated by the hinge helix (23, 26). Conformational heterogeneity is no exception for ScGDH3-K272R (Figure 8, A-D). The cleft opening distance was measured in between the Cα atoms of K119 and F275, for all the subunits of ScGDH3-K272R. We found that the average cleft opening distance is ∼ 16 Å in subunits with “open” conformation but the same is drastically lower in “super-closed” subunits where the average cleft opening distance is ∼ 6 Å (Figure 8D). The difference in cleft opening distance between two set of trimers in the hexameric (biological) assembly is very high (∼ 10 Å) and quite unique for a single GDH structure. The subunit A, B and C (upper trimer) are in “super-closed” conformation and subunit D, E and F (lower trimer) are in “open” conformation. The superimposition of “open” and “super-closed” subunits showed drastic conformational alteration in the hinge helix and overall shift of domain II towards domain I (from “open” to “super-closed”). Our data also supported the fact and showed how subunit conformational alteration occurred upon ligand binding. Apart from the cleft opening distance, major difference among the subunits from different set of trimers can be observed in terms of ligand binding in the active site and at the trimeric interface. Each subunit with “super-closed” conformation has coenzyme (NADP^+^) in the coenzyme binding pocket and citrate in the substrate binding site. However, the “open” subunits do not have any ligand bound to their active sites. We could capture one phosphate molecule at the trimeric interface of “open” subunits. The phosphate group is stabilized by three R399 residues from each subunit (Figure 8E). Surprisingly, the phosphate group is missing at the trimeric interface of the “super-closed” subunits (Figure 8F). All three R399 residues moved away from the trimeric interface. The distances between Cα atoms of each R399 residue were measured and we could observe an increase of ∼ 5 Å at the trimeric interface formed by “super-closed” subunits compared to that of at the trimeric interface formed by “open” subunits. The structural alteration at the trimeric interface was suspected to be directly associated with conformational heterogeneity in ScGDH3-K272R. The presence of both “open” (∼ 16 Å) and “super-closed” (∼ 6 Å) subunit conformational states in single GDH structure is reported for the first time.

**Figure 8:**
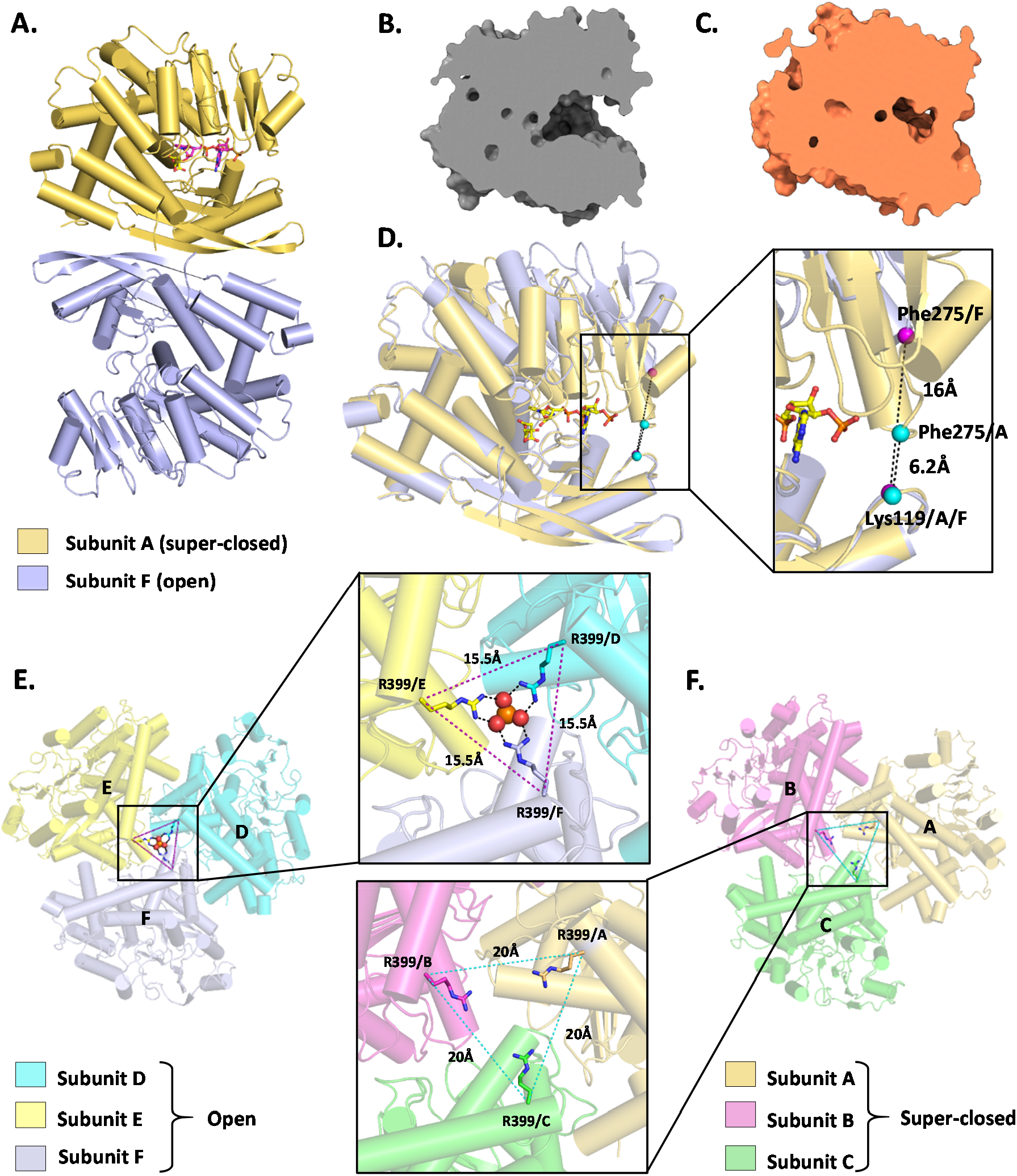
Conformational heterogeneity in ScGDH3-K272R-NADP complex crystal structure. Crystallographic hexameric assembly has been analyzed. **A.** One of the dimeric units showing different conformational states of subunits observed in the crystal structure. The “open” subunit (shown in cartoon, light blue) is in unliganded form and the “super-closed” subunit (shown in cartoon, golden) is complexed with coenzyme, NADP^+^. **B.** Surface clipped representation of the “open” subunit. **C.** Surface clipped representation of the “super-closed” subunit. **D.** Domain I of both these subunits were superimposed to show the significant movement of domain II upon coenzyme (represented in ball and sticks, magenta) binding. Zoomed in view showing the difference in cleft opening between “open” and “super-closed” subunits. The cleft opening distance was measured in between C-alpha atoms (shown in sphere, cyan) of F275 (domain II) and K119 (domain I) for both the subunits and it showed large (∼ 10 Å) difference. **E.** Top view of one set of trimer with subunit D, E, F and all of those are in “open” conformational states. Zoomed in view showing the presence of phosphate ion, stabilized by R399 (black dotted lines represent polar interactions) from each subunit. The C-alpha atoms of R399 of these “open” subunits are 15.5 Å (magenta dotted line) apart from each other. **F.** Top view of another set of trimer with subunit A, B, C and all of them are in “super-closed” conformational states. Zoomed in view is showing the trimeric interface region where the phosphate ion is missing. The C-alpha atoms of R399 (shown in sticks), located in the hinge helix, from each subunit are 20 Å (cyan dotted line) away from each other.

### Binding of NADH to ScGDH3-K272R (MD simulation studies)

The K272R mutant showed significant activity with NADH as coenzyme which is non-specific for the native enzyme. MD simulations were performed in triplicate using K272R complexed with NADH and AKG at the active site. Binding of NADH was studied by comparing with NADP^+^ binding in the crystal structure. The overall binding pose o NADH was found to be slightly different, mainly with respect to the adenine ring (Figure 9A). The adenine ring of NADH was closer to domain II. Placement of ribose moiety, containing 2’OH and other atoms, was similar compared to that of NADP^+^. Further analysis revealed the involvement of highly conserved modified Rossmann fold (β5-α8-β6) and the consensus sequence “GXGXXG/A” (“GSGNVA” in ScGDH3) in supporting NADH binding (Figure 9B). G225, S226, G227, N228 and V229 residues directly interact with NADH in a similar manner, as observed in case of NADP^+^ in the crystal structure (Figure 9C). The involvement of the consensus sequence in identifying and stabilizing the coenzymes [NADP(H) and NAD(H)] inside the binding pocket has been well established (28).

**Figure 9:**
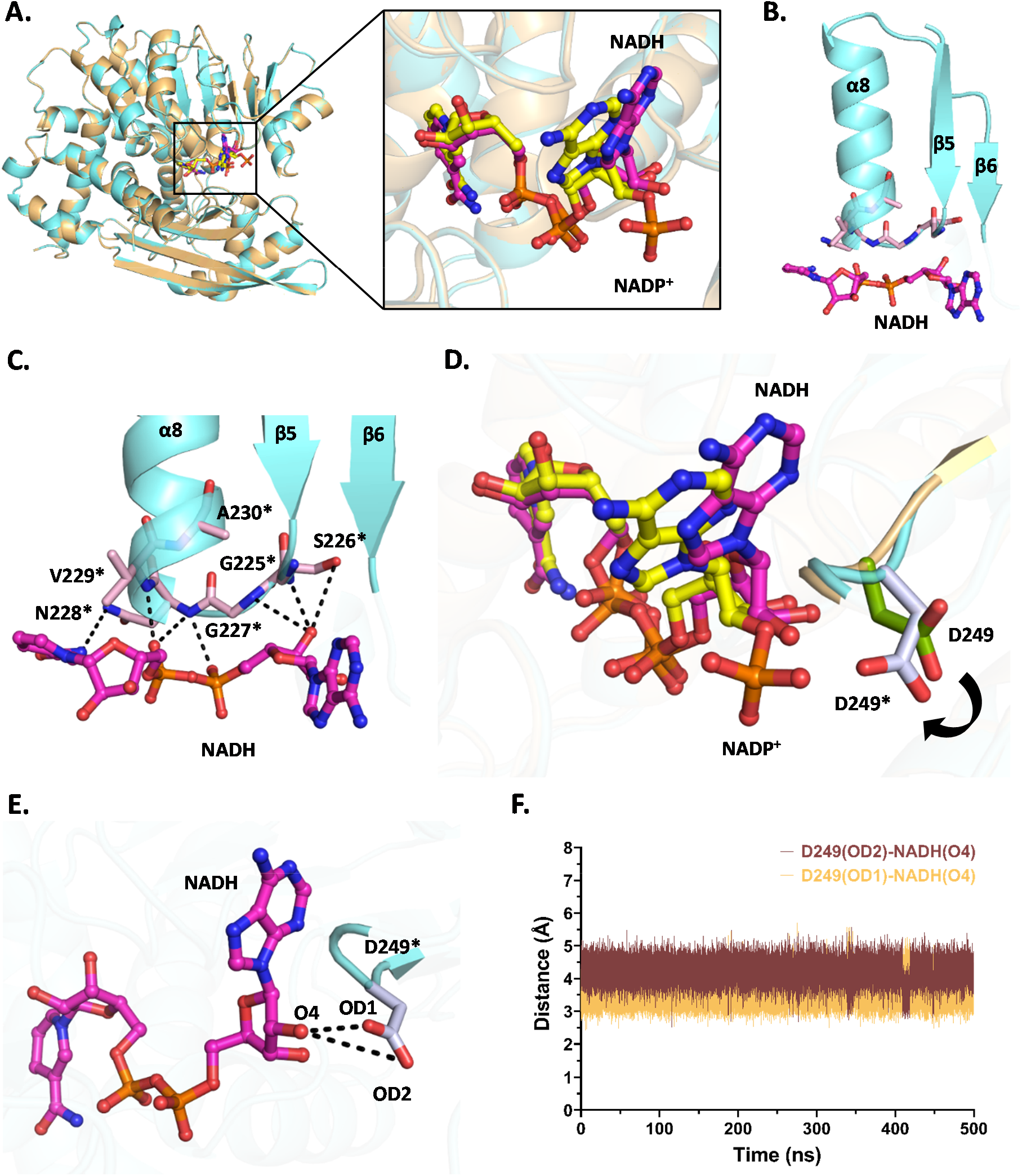
Interactions of NADH with ScGDH3-K272R as observed by the MD simulation studies. **A.** Structural alignment of single subunit from crystal (NADP^+^ bound) and simulated (NADH bound) structures of ScGDH3-K272R. Zoomed in view shows slight alteration in binding poses of NADP^+^ (yellow) and NADH (magenta). Major difference can be observed in the position of adenine ring. Overall binding pose of NADH was closer towards domain II. **B.** The modified Rossmann fold (β-α-β motif) shown in cartoon (cyan), contains the consensus sequence (P1-P6) which helps in binding of NADH inside the coenzyme binding pocket of ScGDH3-K272R. **C.** Polar interactions with G225*-A230* (P1-P6) residues (light pink), part of consensus sequence “GXGXXG/A”, helped in stabilizing the coenzyme inside binding pocket. **D.** Large alteration of D249 side chain was observed NADP^+^ and NADH binding was compared. Side chain of D249 clearly moves towards NADH (denoted by black arrow) while the same remained pointed away from NADP^+^. **E.** Two terminal oxygen atoms (OD1 and OD2) of D249* are showing interaction with O4 atom of NADH. **F.** The trajectories (throughout 500 ns simulation) for interacting distances showing complementary interactions between OD1 and OD2 with the 2’OH (O4) of adenine ribose moiety in NADH. The interaction is maintained throughout the entire simulation. Residues of the simulated structure of the ScGDH3-K272R-NADH complex are marked by “*”. The residues from the ScGDH3-K272R-NADP^+^ crystal structure are not marked.

In spite of the similar binding pose of both the coenzymes, few significant alterations were observed in terms of side chain orientation of the binding pocket lining residues. A significant side chain movement was observed for D249 when NADH is present. The side chain of D249 points away from the coenzyme when NADP^+^ is present and hence it cannot directly interact with the 2’PO_4_ moiety. But in presence of NADH, the side chain moved towards the coenzyme (Figure 9D). Because of this change in side chain orientation, D249 now can directly interact with 2’OH [D249(OD1)-NADH(O4) and D249(OD2)-NADH(O4)] associated with the ribose sugar moiety of NADH (Figure 9E). Little fluctuations were observed in the terminal atoms of D249 (OD1 and OD2) during the simulation. Interestingly, whenever OD1 moves away from the coenzyme and interaction breaks, OD2 comes closer and compensates for the loss of interaction. Hence, the overall interaction was maintained throughout 500 ns simulation (Figure 9F).

The S250 directly interacts with the 2’PO_4_ moiety of NADP^+^. But the side chain moved completely opposite when NADH is present (Figure S6A). Due to this extreme alteration in side chain orientation, S250 is no longer available for interaction with NADH as shown in Figure S6B. Among the residues present in the consensus sequence, S226 showed slightly different side chain orientations when two different coenzyme bindings were considered (Figure S6C). The S226 side chain was pointed towards the NADP^+^ and the interacting distance was 2.7 Å, suggesting a strong polar interaction. However, in case of NADH, the side chain was faced slightly away from the coenzyme. The average interacting distance was observed 3.5 Å-4.0 Å throughout 500 ns simulation, with less than 3.0 Å at several points (Figure S6, D and E). The result suggested higher fluctuations of S226 side chain and weaker interaction compared to that of with NADP^+^.

The R272 residue was involved in strong polar interactions with NADH and helps it to stabilize inside the coenzyme binding pocket. We found three probable interactions that make this mutant (K272R) compatible with NADH as well and aid in carrying out the catalytic reaction. All these three interactions [R272(NH1)-NADH(O4), R272(NH1)-NADH(O8) and R272(NH2-NADH(O12)] were maintained throughout 500 ns simulation with negligible fluctuations. The average interacting distance for all the interactions was 3.5 Å. All the interactions and their trajectories for the complete simulation are shown in Figure 10. The simulation study further explained the molecular basis of how the ScGDH3-K272R mutant becomes catalytically competent with a coenzyme (NADH), non-specific for the native enzyme.

**Figure 10:**
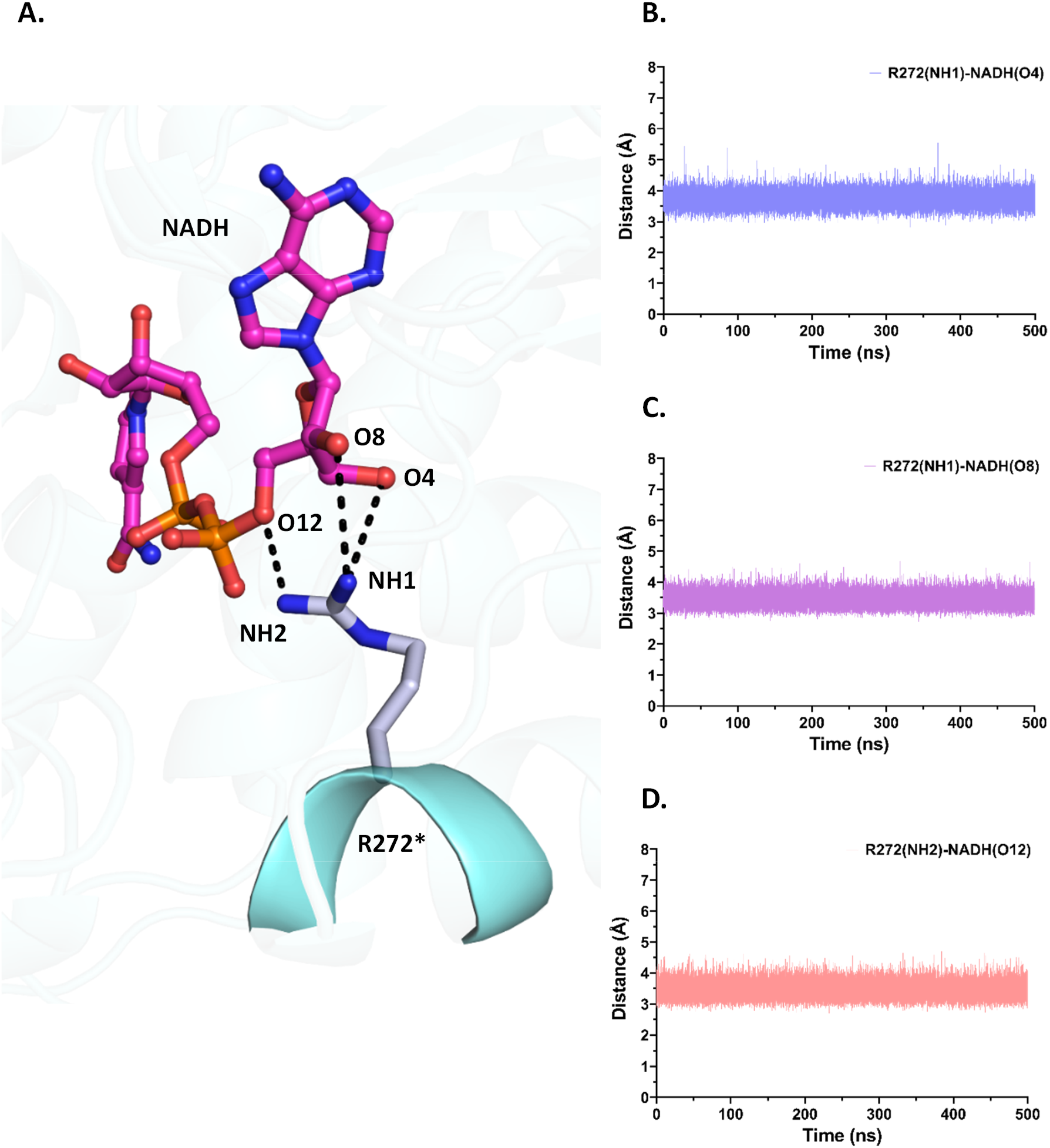
Role of R272 in stabilizing NADH inside coenzyme binding pocket. **A.** R272* (represented in sticks, blue white) shows strong polar interactions with NADH (represented in ball and sticks, magenta). **B.** The plot is showing the trajectory (throughout 500 ns simulation) for interacting distances in between terminal –NH group (NH1) of R272 and 2’OH (O4) of adenine ribose moiety in NADH. **C.** The plot is showing the trajectory for interacting distances in between terminal –NH group (NH1) of R272 and 3’OH (O8) of adenine ribose moiety in NADH. **D.** The plot is showing the trajectory for interacting distances in between terminal – NH group (NH2) of R272 and O12 atom of NADH. Residues of the simulated structure of the ScGDH3-K272R-NADH complex are marked by “*”.

### Evolutionary conservation towards preference of arginine or lysine for coenzyme recognition

As K272R mutation in ScGDH3 exhibited positive effect towards coenzyme specificity, further exhaustive analyses were performed for identifying naturally preferred arginine residue at the same position among the homologous proteins considering a vast and diverse groups of oxidoreductase including NAD-specific GDHs, NADP-specific GDHs and other amino acid dehydrogenases (leucine dehydrogenase, phenylalanine dehydrogenase) from the organisms across different kingdoms. The analyses were executed with a primary focus on the structural alignment instead of sequence alignment as later often produced ambiguous results for GDHs. To our surprise, the presence of a structurally homologous arginine was identified in some of the well characterized NAD-specific GDHs: *Clostridium symbiosum* GDH (CsGDH, PDB ID: 1AUP, 29), *Bacillus subtilis* GDH (BsGDH, PDB ID: 3K8Z, 30) and *Lysinibacillus composti* GDH (LcGDH, PDB ID: 8I83, 31). Structure-based sequence alignment of ScGDH3-K272R-NADP^+^ complex (PDB ID: 47UL) with well characterized NADP-specific GDH (PDB IDs: 5XVX) and NAD-specific GDHs (PDB IDs: 1AUP, 3K8Z, 8I83) shows that presence of lysine and arginine respectively, at this specific position is conserved (Figure S7A). Domain II based alignment of AnGDH, ScGDH3-K272R and CsGDH structures resulted in perfect alignment of the modified Rossmann fold and the helix containing concerned K/R residues (Figure S7B). The alignment also clearly showed the arginine (in CsGDH and ScGDH3-K272R mutant) is occupying more space inside the coenzyme binding pocket than that of the lysine (in AnGDH) (Figure S7C). Separate structural alignment among the mentioned NAD-specific GDHs revealed the position and orientation of the arginine side chain to be similar although the structural fold is slightly different (Figure S7, D and E). Among other amino acid dehydrogenases, one NAD-specific phenylalanine dehydrogenase from *Rhodococcus sp.* has an arginine (R210) at the homologous position. The NAD^+^-complexed crystal structure (PDB ID: 1C1X) of this enzyme showed R210 is directly interacting with the 3’OH of adenine ribose moiety of NAD^+^ to stabilize the coenzyme inside the binding pocket (32). Similarly, the presence of homologous arginine was identified in an NAD-specific leucine dehydrogenase from *Pseudomonas aeruginosa* (PDB ID: 7VID). Although it is an apo structure but the similar orientation of the R202 side chain inside the coenzyme binding pocket suggested possible interaction with NAD(H) (33). These analyses indicate that presence of an arginine is also preferred by nature for imparting NAD-specificity in amino acid dehydrogenases.

## Discussion

GDH is an essential metabolic enzyme which plays a critical regulatory role in all living organisms. This regulation occurs through layers of interconnected events inside the cells in order to maintain the continuity of both the metabolic cycles at any point of time (2, 3, 4). One of the important parts of this regulation takes place via coenzyme selection. It has been very well established that the directionality of the catalytic reaction highly depends on which coenzyme is being used (9). The importance of modulation of size of the coenzyme binding pocket to accommodate appropriate coenzyme based on the physiological need, has been explained in case of dual coenzyme specific mammalian GDHs (14). However, to understand the molecular basis of the coenzyme selection, fungal GDHs are quite unexplored. This motivated us to proceed with understanding the molecular basis of coenzyme preference and coenzyme binding in *S. cerevisiae* NADP-specific GDH3 (ScGDH3).

This study presents mutational studies on ScGDH3 guided by evolutionary conservation analysis, their extensive biochemical characterizations and structural elucidation to explain altered coenzyme specificity, coenzyme binding, catalytic efficiency and the overall enzyme functional regulations.

### Alteration in biochemical properties of the mutants

In this present study, we have considered the size/volume of coenzyme binding pocket as one of the critical parameters to be responsible for coenzyme selection in fungal GDHs. We hypothesized reduced pocket volume with residues interacting with the 2’PO^4^ moiety might facilitate NAD(H) binding. The single (K272R) mutation aided in improved selection of NADH and NAD^+^ which are non-specific coenzymes for the native enzyme, followed by tremendous improvement in the affinity and catalytic efficiency. This is indeed a unique observation which proves our hypothesis to be true. Coenzyme binding study showed K272R mutant exhibits significant improvement in binding affinity towards NAD(P)H. Here, it was possible to work with a broad range of coenzyme concentration (0.005-250 μM) unlike in kinetic assays (1-250 μM) due to limited instrument sensitivity. The K272R mutant significantly alters several important biochemical and kinetic properties present in the WT enzyme. The molecular basis of this alteration is discussed below.

The S250E mutant also showed introduced dual-coenzyme specificity, but this time it is more stringent compared to the K272R. It appears that strong repulsion from the negatively charged carboxylate group of the mutated E250 destabilized NADP(H) in the binding pocket and resulted in drastic decrease in enzyme activity. The same residue might have facilitated favourable interactions with non-specific coenzymes [NAD(H)] and which results in enzyme catalysis with higher efficiency as compared to the native enzyme. The S250E mutant has higher affinity towards NADH than that of NADPH. Hence, it is evident that the S250E mutation modulated the binding pocket microenvironment in such a way that the preference for NADH binding by the mutant is higher than the specific coenzyme (NADPH); and thus indicating the importance of the S250 residue in determining coenzyme specificity in NADP-specific GDHs.

The contribution of domain I in coenzyme recognition by GDH was assessed. Notably, two single mutants (K119F and K119A) K119 individually showed drastic reduction in the specific activity. The mutants exhibited significantly less NADP(H)-specific activity as compared to the native enzyme and almost no NAD(H)-specific activity. Although domain I of small hexameric GDHs is known not to contribute in coenzyme specificity (18, 21), our study demonstrated domain I residue plays a crucial role in regulating NADP-specific catalytic activity in ScGDH3.

### Structural insights and conformational flexibility

Our study presents the first high resolution structure of GDH from *Saccharomyces cerevisiae*. Structural analysis provided many important and interesting insights into the coenzyme selection, coenzyme binding and conformational heterogeneity.

In the hexameric structure, only half of the subunits (one set of trimer) is complexed with coenzyme NADP^+^ and in “super-closed” conformation. In the active site of these “super-closed” subunits, citrate was captured instead of the actual substrate, AKG. We assume the crystallization mother liquor was the source of citrate present in the active site. Although AKG (4 mM) was used for co-crystallization, it was absent in the substrate binding site of any of the subunits. Two practical possibilities can be figured out: first possibility, the substrate binding site might be occupied by citrate which was present in much higher concentration (1.8 M) than that of AKG (4 mM). Capturing carboxylate group containing compounds, present in the mother liquor have also been reported for *Aspergillus terreus* GDH (AtGDH) by our group (27). Second possibility, the catalytic reaction might have occurred as the mother liquor had ammonium (NH_4_^+^) ion, necessary for the reaction at a suitable pH. For co-crystallization, we used 10-fold higher concentration of both the substrate and coenzyme than that of the enzyme. It is possible, that all AKG molecules might have been consumed during the crystallization process. The presence of NADP^+^ also supports the possibility of complete utilization of the substrate.

GDH is an intrinsically dynamic enzyme; it exhibits flexible domain movement and different subunit conformational states as parts of the functional regulation (23, 26). GDH3 from *Saccharomyces cerevisiae* (ScGDH3) is no exception in this aspect. Our structure showed two distinct subunit conformations: “open” and “super-closed”. Previous studies shown that only NADPH is sufficient in initiating domain II movement and subsequent closing of the cleft (23). This observation was further supported by our structure. The “super-closed” conformation helps to accommodate the coenzyme by forming the binding pocket. It also aids in interaction with domain I residues (R79, H81, K119 and D151). The R399 residue located in the hinge helix (α_14_) which regulates the extent of domain II movement during conformational alteration. Due to huge domain dynamicity, R399 moved away from the trimeric interface and no longer available for stabilizing the phosphate ion. This could be the possible reason for missing phosphate ion at the trimeric interface of “super-closed” subunits. The presence of “open” and “super-closed” subunits in a single structure is unique and provides clear evidence of extreme conformational flexibility of this enzyme. Our study reports this unique oligomeric assembly for the first time which is captured using X-ray crystallography.

### Molecular insights into the altered coenzyme specificity

The crystal structure of ScGDH3-K272R complexed with NADP^+^ revealed the interactions in which the mutated R272 is involved for binding of the coenzyme. The K272 in the WT enzyme is highly conserved among all other NADP-specific GDHs. Several coenzyme (NADPH) bound structures of GDHs from other organisms showed conserved interactions as well (23, 24). In our structure, mutated R272 is involved in interactions, particularly with NADP^+^. Both the terminal amino groups (NH1 and NH2) of the residue are involved in strong polar interactions with the 2’PO_4_ moiety of the coenzyme. Bigger side chain of R272 occupied more space in the coenzyme binding pocket and strictly specific interactions only with the coenzyme increase the affinity (*1/K_M_*) towards NAD(P)H for ScGDH3-K272R as compared to the native enzyme. More stringent interactions have also increased the coenzyme binding affinities for K272R mutant as compared to the native enzyme.

MD simulation studies showed how the microenvironment inside coenzyme binding pocket aided in not only appropriate binding of NADH but also becoming catalytically competent with this non-specific coenzyme. Significant alterations can be observed in terms of side chain orientation of some important residues (discussed in the result section) favoured NADH binding and stabilization. Crystal structure of ScGDH3-K272R complexed with NADP^+^ has explicitly explained the molecular basis of tremendous increase in coenzyme affinity achieved by mutating a single amino acid residue (K272R). Our MD simulation studies further showed that the R272 residue is highly capable of stabilizing NADH inside the coenzyme binding pocket. Both our crystal structure and simulation study further established the important role of extremely conserved P7 residue (D249 in ScGDH3) in recognizing the appropriate coenzyme. The side chain of this negatively charged residue remained away from the 2’PO_4_ moiety of NADP^+^ to avoid any electrostatic repulsion that destabilize the coenzyme inside the binding pocket. The same residue oriented towards NADH and interacted via hydrogen bond formation with the 2’OH associated with the adenine ribose, to stabilize the coenzyme. Similar behaviour of the P7 residue upon binding of these two coenzymes, was reported by Smith and his group in case of a bovine (*Bos taurus*) GDH (BtGDH) which shows dual coenzyme specificity. This further strongly supported our observation that ScGDH3-K272R mutant truly exhibits dual coenzyme specificity.

### Importance of K272R mutation in the light of evolution

The lysine (K272 in ScGDH3) is highly conserved among NADP-specific fungal GDHs and facilitates binding of the 2’PO_4_ moiety of NADPH. Overall, our findings indicated the presence of a homologous arginine in several naturally occurring NAD-specific dehydrogenases including GDHs. The position of the arginine side chain located deep inside the coenzyme binding pocket and its strong interaction with the adenine ribose moiety of NAD(H) exactly match with our MD simulations results. All these observations further explain how ScGDH3-K272R mutant gained NAD-specificity as ScGDH3-WT is a natively NADP-specific GDH.

Considering NADP-specific GDHs, lysine is more or less conserved in that position across different species. However, NADP-specific GDH from *Corynebacterium glutamicum* (CgGDH) highlighted an interesting aspect. The NADP^+^-complexed crystal structure (PDB ID: 5IJZ) showed an arginine (R290) is facing towards 2’PO_4_ moiety of NADP^+^ and interacting with it (22) (Figure S7F). The conserved lysine (K284) is present but not interacting with the 2’PO_4_ moiety, which is supposed to interact. Hence, it is evident that arginine can be crucial in stabilizing NADP(H) as well inside the coenzyme binding pocket, which also sheds light on the increased NADPH affinity in ScGDH3-K272R mutant.

All these facts strengthen our experimental findings and again proved our hypothesis to be accurate. Our global rational approach towards mutational studies, biochemical findings and their appropriate structural explanation would pave the way for understanding coenzyme selection process in fungal NADP-specific GDHs.

## Conclusions

Our study provides a holistic overview of the molecular basis of coenzyme selection in NADP-specific ScGDH3 and other fungal GDHs. Single point mutation (K272R) results in altered coenzyme specificity with drastic increase in affinity (*1/K_M_*), catalytic efficiency and binding affinity (*1/k_d_*) for both the coenzymes. Another point mutation (S250E) introduced dual coenzyme specificity with comparable NAD(P)H affinities (*1/K_M_*). The data successfully established the role of domain I residue (K119) in regulating coenzyme specific catalytic activity of ScGDH3. We also solved the first NADP^+^-complexed structure of GDH mutant (K272R) from *Saccharomyces cerevisiae*, in which we could successfully captured two distinct subunit conformations: “open” (without ligands) and “super-closed” (with ligands). Crystal structure analysis deciphered the molecular basis of increased NADPH affinity in ScGDH3-K272R. Results from our MD simulation studies further provided crucial molecular insights into the dual specificity of ScGDH3-K272R and explained how NADH gets stabilized inside the coenzyme binding pocket. Both our biochemical and structural findings helped to establish the process of coenzyme binding. All these residues are highly conserved among other NADP-specific GDHs. Hence, findings of this study would indeed be applicable to understand enzyme functional regulation via coenzyme selection in other NADP-specific GDHs across living organisms. The rational approach behind the mutagenesis study by considering coenzyme binding pocket size can be applicable for other dehydrogenases as well.

## Experimental procedures

### Cloning and overexpression of wild type ScGDH3 (ScGDH3-WT)

*Scgdh3* gene was amplified from the *S. cerevisiae* genomic DNA (obtained from Prof. P. J. Bhat, Department of Biosciences and Bioengineering, IIT Bombay) and inserted in between *NdeI* and *XhoI* restriction sites of pET28a(+) vector with an N-terminus His_6_-tag. The expression construct also has a Tobacco Etch Virus (TEV) protease cleavage site in between the His_6_-tag and *Scgdh3* gene to facilitate removal of the tag after the purification via affinity chromatography. The successful construction of the ScGDH3-WT expression vector was confirmed by sequencing.

The recombinant plasmid with *Scgdh3* gene was transformed into ΔGDH *E. coli* BL21(DE3) expression cells. Single colony was grown overnight as a primary culture in LB (Luria-Bertani) broth supplemented with kanamycin (50 µg/ml) at 37 °C. 1% (v/v) of primary inoculum was added into 50 ml LB broth supplemented with kanamycin (50 µg/ml) and grown in a shaking incubator at 37 °C until 1.3-1.5 optical density (at 600 nm). Next, the cells were treated with 1000 μM isopropyl 1-thio-β-D-galactopyranoside (IPTG) and overexpression of ScGDH3 was carried out at 30 °C for 5 hours by shaking the culture at 100 rpm. The cells were harvested at room temperature by centrifugation (Multifuge X3R, Thermo Scientific) at 8000 rpm for 10 min. The cell pellet was then resuspended in a lysis buffer [100 mM Tris-Cl buffer, pH 7.5, 1 mM phenylmethylsulfonyl fluoride (PMSF)]. The cell suspension was homogenised by ultrasonication at pulse 2 sec on-off and 40% amplitude. The cell lysate was then centrifuged at 4 °C (Multifuge X3R, Thermo Scientific) at 16,000 g for 30 mins. The protein content in the supernatant and pellet was analysed using 12% SDS-PAGE (34). The protein concentration was estimated using Bradford’s method (35). All the steps starting from cell lysis were performed at 4 °C.

### Purification of ScGDH3-WT

For the purification of ScGDH3-WT, Ni-NTA affinity chromatography purification was performed using FPLC (ÄKTA pure, Cytiva) as the first step. A pre-packed HisTrap^TM^ Fast Flow (FF) 5 ml Ni-NTA column was equilibrated with the binding buffer 1 (BB1) (100 mM Tris-Cl buffer, pH 7.5). After homogenisation and centrifugation, cell free extract was filtered (using PVDF membrane, 0.45 µm) and loaded on to the column at 1 ml/min flow rate, followed by washing using 5% (25 mM imidazole) and 10% (50 mM imidazole) elution buffer (EB) (100 mM Tris-Cl buffer, 500 mM imidazole, pH 7.5) to remove the unbound proteins. The His_6_ tagged proteins were eluted with 20% (100 mM imidazole) and 50% (250 mM imidazole) elution buffer. The eluted fractions were pooled together, the combined sample was subjected to buffer-exchange to get rid of imidazole and concentrated (up to ∼ 2 ml) using Amicon ultrafiltration concentrator (∼ 50 kDA cut off). The total protein amount was estimated using Bradford’s method. The final step of purification was performed using a Superdex 200pg 16/600 size-exclusion column (Cytiva), equilibrated with the binding buffer 2 (BB2) (20 mM Tris-Cl buffer, pH 7.2). Protein was loaded at 0.5 ml/min and followed by the elution. The eluted fractions were collected and analyzed on 12% SDS-PAGE (34) to check protein purity. The pure ScGDH3-WT fractions were pooled and used for further studies.

### Site-directed mutagenesis, expression and purification of the mutants

Fungal GDHs have two different coenzyme specificities; either NADP(H)-specific or NAD(H)-specific. Although the coenzymes have remarkably similar molecular structures, the specificity for these coenzymes is very stringent among fungal GDHs. The major difference between these two coenzymes is the presence of a phosphate (PO_4_^3-^) group associated with the nicotinamide moiety of NADPH. Hence, to understand the molecular basis of coenzyme recognition, we particularly focused on those residues which directly or indirectly interact with the 2’PO_4_ moiety. We found four such residues: K119, D249, S250 and K272; all these residues are extremely conserved among all the other NADP-specific fungal GDHs (Figure S8). Their interactions can be visualized from a high-resolution coenzyme-bound crystal structure (PDB ID: 5XVX) of *A. niger* GDH, solved by our group (23) (Figure S9). Among these residues, D249 is the very important P7 (acidic) residue which is conserved among all the dehydrogenases including GDHs. K119 is the only residue from domain I that directly interacts with the 2’PO_4_ moiety. Hence, these residues were selected for mutational studies in ScGDH3. Based on the nature of interaction with the 2’PO_4_ moiety, types of respective amino acids and sequence alignment (with NAD-specific GDHs) analysis, several point mutations were proposed initially: K272R, K272A, S250E, K119A, K119F. One of the major rationale behind proposing these point mutations was to reduce the overall size of the coenzyme binding pocket so that NADH can easily accommodate inside it. Our aim was to check whether mutating these residues would impart any altered coenzyme specificity in a natively NADP-specific enzyme. Arginine (R) has bigger side chain than lysine (K); it can easily mimic the interactions that K272 was involved in and can stabilize NADH inside the binding pocket at the same time. So, K272R mutation was proposed. Serine (S) was mutated to glutamate (E) as it has longer side chain with overall negative charge, which can destabilize NADPH by repelling 2’PO_4_ moiety and stabilize NADH by forming hydrogen bond with the 2’OH group. Based on the rationale mentioned above, ScGDH3-K272R-S250E was generated and the cumulative effect in the biochemical properties of ScGDH3 was assessed.

Recombinant plasmid (pET28a(+)) containing the Sc*gdh3* gene was used as template for Polymerase Chain Reaction (PCR). Specific sets of primers were synthesized by Sigma-Aldrich^®^ to favour the mutation at the specific site of the gene. Sequences of all the primers used for mutagenesis studies are listed in the table (Table S1) below. The reaction mixture (50 μl) comprised of template DNA (1 μl), forward and reverse primers (10 μM, 2.5 μl each), 5X Q5 reaction buffer (10 μl), dNTPs (10 mM, 2 μl), Q5 DNA polymerase (NEB) (0.5 μl) and nuclease-free water (31.5 μl). The reaction condition includes preheating at 98 °C for 10 min, 16 cycles of initial denaturation (at 98 °C for 40 s), annealing (at 68 °C for 40 s), extension (at 72 °C for 10 min); and final extension at 72 °C for 10 min. PCR amplified products were subjected to DpnI (NEB) digestion at 37 °C for 2 hours. The digested products were cleaned using PCR-clean up kit (Qiagen) and transformed into CaCl_2_ treated competent *E. coli* DH5α cells by heat shock at 42 °C for 90 sec and immediate cooling in ice. The transformed cells were screened by growing them overnight on kanamycin (50 µg/ml) supplemented Luria agar (LA) plates. The transformed colonies were chosen randomly and grown at 37 °C in kanamycin (50 µg/ml) supplemented LB broth. The plasmid was isolated using Exprep^TM^ Plasmid SV mini (GeneAll) and concentrated up to ∼ 80 ng/µl. All the point mutations were confirmed by DNA sequencing. Isolated plasmids were transformed into CaCl_2_ treated competent ΔGDH *E. coli* BL21(DE3) cells.

Expression of all the ScGDH3 mutants were performed following the exact protocol described for ScGDH3-WT except ScGDH3-K119A and ScGDH3-K272R-S250E which showed extremely less soluble expression. Both of these mutants were overexpressed following a different protocol altogether. Cells were grown at 37 °C until 0.2-0.3 optical density (at 600 nm). Cells were then kept at 16 °C for 45 minutes without any shaking. Next, the cells were treated with 200 μM and 50 μM IPTG respectively and induction was carried out at 16 °C for 20 hours by shaking at 100 rpm. Purification protocol for all the mutants was exactly same as followed for ScGDH3-WT. Overall structural folding of these mutants was assessed by performing circular dichroism (CD) experiment.

### Enzyme activity assays

The enzyme activity assay was performed based on the previously established method (36). Forward activity of NADP-specific glutamate dehydrogenase is considered as the reductive amination of α-ketoglutarate (AKG) to form L-glutamate in presence of excess ammonia along with the oxidation of NAD(P)H to NAD(P). The hydride (H^-^) transfer was spectrophotometrically monitored at 340 nm (A_340nm_) to measure the enzyme activity. The assay was performed at room temperature, using 1 ml reaction mixture contains assay cocktail (50 mM Tris-Cl, pH 7.2, 8 mM AKG, 50 mM NH_4_Cl), 0.2 mM NAD(P)H and purified enzyme (dilution required to get 5-10% conversion of NAD(P)H). The change in absorbance (rate of disappearance of NAD(P)H) was monitored in UV/Vis Spectrophotometer (JASCO, Japan) over a period of 60 s and ΔA_340nm_ was calculated. One enzyme activity unit corresponds to the amount of enzyme required to oxidize one µmole of NAD(P)H per minute at standard assay conditions.

The reverse enzyme activity is considered as the oxidative deamination of L-glutamate to form AKG along with the reduction of NAD(P) to NAD(P)H. This assay was also performed at room temperature but using a different setup. The assay cocktail contains assay buffer (100 mM glycine-NaOH, pH 9.5) and 50 mM L-glutamate. Additionally, 2 mM NADP^+^ and 5 mM NAD^+^ (individual assay) were added with purified enzyme to prepare a reaction mixture of 1 ml. The rate of appearance of NAD(P)H was spectrophotometrically monitored at 340 nm (A_340nm_) for 60 s to measure the enzyme activity. One enzyme activity unit corresponds to the amount of enzyme required to reduce one µmole of NAD(P)^+^ per minute at standard assay conditions.

### Enzyme kinetics of ScGDH3-WT and mutants

Coenzyme saturation kinetics were performed for ScGDH3-WT and mutants (K272R, S250E) based on both reductive amination (forward) and oxidative deamination (reverse) reactions as described before (36). For forward reaction, coenzyme (NAD(P)H) concentration was varied (1-250 μM) while AKG and NH_4_Cl were used at their saturating concentration of 8 mM and 50 mM, respectively. Enzyme kinetics was performed at room temperature, following the standard assay protocol.

For reverse reaction, varied concentrations of NADP (0.0005-3 mM) and NAD (0.5-20 mM) were used while L-glutamate was used at its saturating concentration of 50 mM. Enzyme kinetics was performed at room temperature, following the standard assay protocol. All the kinetics assays were performed in triplicates. The SigmaPlot 12.3 and GraphPad Prism 8.4.2 were used for plotting and analysing the data.

### Tryptophan fluorescence quenching assays

Purified ScGDH3 samples (WT and mutants) in 20 mM Tris-Cl buffer, pH 7.2, were used for tryptophan fluorescence quenching assay to assess qualitative binding of coenzymes. The change in fluorescence intensities in the presence of NAD(P)H were recorded. The excitation wavelength was 285 nm and the emission wavelength was recorded in the 300-500 nm range using the spectrofluorimeter (JASCO, Japan). Both the band width and slit width were kept at 2.5 nm. The detector sensitivity was optimized and set as medium. Total reaction mixture volume was 500 μl which contained a fixed concentration (6 μM) of enzyme (WT and mutants), and varying concentrations (10-250 μM) of NADPH/NADH. The inner filter effect was corrected from the observed fluorescence intensities. All the measurements were performed in triplicates at the room temperature. The GraphPad Prism 8.4.2 software were used for plotting and analysing the data.

### MicroScale Thermophoresis (MST)

Both the enzymes (ScGDH3-WT and ScGDH3-K272R) were labelled with His-Tag Labelling Kit (RED-tris-NTA 2nd Generation, NanoTemper Technologies) which directly interacts with the His-tag, present in these proteins. The enzyme and fluorophore were mixed in equal proportion in terms of their molar concentrations (200 nM) and incubated for 30 min at 4 ℃. The mixture was centrifuged at 10000 rpm for 10 min at 4 °C to remove the unbound fluorophore. The labelling was confirmed by measuring the fluorescence counts at 650 nm and 670 nm. Ligand binding assays were performed using 16 samples with different coenzyme concentrations (0.003-250 µM). These samples were prepared by serial dilution method. Initially, 10 µl buffer (20 mM Tris-Cl, pH 7.2) supplemented with 0.01% Tween-20 was added to each 16 tubes except the first one. Then 20 µl of coenzyme [NAD(P)H] (500 µM) was added into the first tube, 10 µl of which was aspirated, added and mixed into the 2^nd^ tube. Similarly, coenzyme was serially diluted up to the 16^th^ tube. Finally, 10 µl of fluorophore labelled enzyme was added and mixed in each tube. Hence, the final concentration of labelled enzyme in each tube was 100 nM. All the samples were then incubated at room temperature for 30 min and each sample was aspirated using monolith premium capillary tubes (NanoTemper Technologies). The binding assay was performed using the Monolith X instrument (NanoTemper Technologies. The binding affinity was measured by monitoring ligand-induced spectral shifts in the fluorescence emission of the labelled enzyme. In the dose response curve, the ratio of fluorescence intensities at 670 nm and 650 nm was plotted as a function of ligand (coenzyme) concentration using SigmaPlot 8.4.2 software. Dissociation constant (*k_d_*) was determined based on the spectral shift analysis using NanoTemper software.

### Crystallization

ScGDH3-K272R was used for crystallization following both sitting and hanging drop vapour diffusion methods. Highly pure and concentrated enzyme combined with the substrate (AKG) and both the coenzymes (NADPH/NADH) were used for setting up the initial crystallization screen. Molar concentrations of enzyme, substrate and coenzymes were 400 µM (20 mg/ml), 4 mM and 4 mM respectively, in the protein preparation used for the crystallization. The ternary complex was incubated for 30 min at room temperature before setting up the crystallization drops. Initial screens were set up in 96 well Intelliplate (Art Robins) using commercial crystallization screen solutions [PEG Suite (Molecular Dimensions), PEG Rx (Hampton Research), PEG/Ion (Hampton Research) and Index (Hampton Research)] with the help of Phoenix (Art Robins) robot (Protein Crystallography Facility, IIT Bombay). The crystallization trays were incubated in a vibration-free crystallization incubator at 18 °C and frequently observed to assess the growth of crystal(s) under the stereomicroscope. Upon getting several hits, hanging drops were set up using the ternary complex with varying the drop size and different reagent concentrations to obtain bigger and good quality crystals. The best hit was obtained in the crystallization screening solution containing 1.8 M Ammonium citrate tribasic at pH 7.0 (INDEX, Hampton Research) and it grew maximum in size within 30 days.

### X-ray diffraction data collection and processing

X-ray diffraction datasets were collected by the rotation method. Cryoprotectant solution was prepared by adding 25% (v/v) glycerol in the corresponding mother liquor to prevent ice formation during crystal freezing. The crystal was quickly transferred to the cryoprotectant solution and subsequently flash-frozen at 100 K in the liquid nitrogen stream. The X-ray diffraction dataset for the ScGDH3-K272R-ternary complex was collected by rotation method with an oscillation of 0.5° per frame at the Indus-2 synchrotron beamline (PX-BL21) (37) of Raja Ramanna Centre for Advanced Technology (RRCAT), Indore, India. Dataset was indexed, integrated and scaled using XDS (38). F2MTZ and CAD in CCP4 (39) were used to convert the intensities into structure factors. Table 5 presents the data collection statistics.

### Structure determination, model building and refinement

The structure of the ScGDH3-K272R complexed with NADP^+^ was determined by molecular replacement method. The structure of *Aspergillus niger* GDH (AnGDH, PDB ID: 5XVX) was used as the template for the molecular replacement. Matthew’s coefficient (40) value of 3.09 Å^3^ Da^-1^ indicated the presence of four molecules in the asymmetric unit. Domain I and domain II of the template were separated and used as two different templates. Then both the templates were provided simultaneously to PHASER (41) for the search. Initially, both the templates were used to search all the domains of four molecules. All the domains were correctly placed by PHASER. Then the domains were merged and the correct orientations of four subunits were generated. After phasing, manual model building was performed using COOT (42). The model was then subjected to iterative refinements using REFMAC5 in CCP4 (43) and phenix.refine in PHENIX (44). Among four subunits, two subunits were in “open” conformation and remaining two exhibited “super-closed” conformation. Based on further analysis of the *F_o_-F_c_* electron density map, citrate and NADP^+^ were modelled in the substrate binding site and coenzyme binding pocket respectively, of the “super-closed” subunits. These ligand densities were missing in the subunits with “open” conformation. The ions and solvent molecules were gradually added while iterative refinements were performed. The improvement of the overall structure quality (figure of merit, FOM) and decrease of *R*_work_/*R*_free_ were continuously monitored. Table 5 presents the refinement statistics.

### Molecular Dynamics (MD) simulation studies

Our aim was to investigate the alteration in the binding pocket and interactions of residues with NAD(H). We tried to obtain a crystal structure of NADH bound to ScGDH3-K272R, but the complex didn’t crystallize. Hence, MD simulation was performed using ScGDH3-K272R complexed with NADH to compare any alteration in the coenzyme binding pocket which might be useful in explaining the molecular basis of changes in the biochemical properties. We started with our crystal structure (PDB ID: 47UL) where NADH was positioned inside the coenzyme binding pocket by deleting 2’PO_4_ moiety of NADP^+^. The NADH binding pose was optimized using crystal structure (PDB ID: 1AUP) of *Clostridium symbiosum* GDH (CsGDH) complexed with NADH. The substrate, AKG was also placed at the substrate binding site following the coordinates of AKG present in AnGDH-ternary complex structure (PDB ID: 5XVX).

All the simulation studies were carried out with GROMACS v2022.2 (45) using CHARMM36 force field (46). Protein–ligand complexes were positioned in a cubic box of water molecules in the macromolecular simulation system, using the CHARMM-GUI server (47). The protein and the box’s borders were kept at least 10 Å apart. To reach the required physiological ionic strength of 0.15 mol·L^-1^, the accurate number of Na^+^ or Cl^-^were added (ion placing method: Monte-Carlo) and system was neutralized. In order to eliminate any near contacts or steric clashes, the system was then energy-minimized. NVT (constant number of particles, volume, and temperature) and NPT (constant number of particles, pressure, and temperature) ensembles were among the steps of equilibration that were applied to the system for a total duration of 5 ns. Positional restraints were applied to the protein backbone, side chains, and bound ligands (NADH and AKG) during equilibration and production md, in order to maintain molecular architecture of the active site while providing flexibility to the solvent environment. Cα backbone and side chain atoms were restrained using force constants of 1000 kJ mol^-1^ nm^-2^ and 200 kJ mol^-1^ nm^-2^, respectively. The modified Berendsen velocity-rescaling thermostat (v-rescale) (48) regulated the temperature at 303.15 K during equilibration, while the Parrinello-Rahman barostat (49) regulated the pressure. The Particle Mesh Ewald (PME) method (50) was utilized to compute the long-range electrostatic interactions and non-bonded interactions with a real-space cut-off of 1.2 nm. After equilibration, the production MD simulations were performed for 500 ns in triplicate. During the total time of 1.5 μs simulation, atomic coordinates were captured every 6.67 ps. Visualization of trajectories were performed using VMD and graphs were plotted using Graph Pad Prism 8.4.2.

## Data availability

The atomic coordinates of ScGDH3-K272R-NADP complex was deposited in the Protein Data Bank (PDB) with accession code 47UL. All relevant data are available from the corresponding author upon request.

## Supporting information

This article contains a supporting information.

## Acknowledgements

We thank Prof. Paike Jayadeva Bhat for providing *S. cerevisiae* genomic DNA and Prof. Narayan S. Punekar for providing ΔGDH *E. coli* BL21(DE3) expression strain. We thank Dr. Ravindra D. Makde, Dr. Ashwani Kumar and Dr. Biplab Ghosh for their kind support in diffraction data collection at Indus-2 synchrotron beamline (PX-BL21) of Raja Ramanna Centre for Advanced Technology (RRCAT), Indore, India. We acknowledge the “Protein Crystallography Facility” and “MST Facility” at IIT Bombay. We acknowledge partial financial support from Science and Engineering Research Board (SERB) (Grant No. CRG/2021/002404), and Anusandhan National Research Foundation (ANRF) (Grant No. ANRF/ARG/2025/001341/LS), Govt. of India.

## Author contributions

A. B., A. R., S. S., B. K. J. G., and P. B. writing–review & editing; A. B. and P. B. writing– original draft; A. B., A. R., and P. B. visualization; A. B., A. R., and P. B. validation; A. B., A. R., S. S., and B. K. J. G., methodology; A. B., B. K. J. G., and P. B. investigation; A. B., A. R., S. S., B. K. J. G., and P. B. formal analysis; A. B., A.R., and S. S. data curation; P. B. conceptualization; P. B. supervision.

## Funding information

The work was partly supported by research funding to P. B. from Science and Engineering Research Board (SERB) (Grant No. CRG/2021/002404), and Anusandhan National Research Foundation (ANRF) (Grant No. ANRF/ARG/2025/001341/LS), Govt. of India.

## Conflict of interest

The authors declare no competing interests with the contents of the article.

## Abbreviations and nomenclature

GDH: glutamate dehydrogenase
NADPH: nicotinamide adenine dinucleotide phosphate, reduced form
NADP^+^: nicotinamide adenine dinucleotide phosphate, oxidized form
NADH: nicotinamide adenine dinucleotide, reduced form
NAD^+^: nicotinamide adenine dinucleotide, oxidized form
MD: molecular dynamics
AKG: αketoglutarate
TCA: tricarboxylic acid
WT: wild type
MST: microscale thermophoresis
PAGE: poly acrylamide gel electrophoresis
PCR: polymerase chain reaction

## Notes

### Competing Interest Statement

The authors have declared no competing interest.

