## Supporting information for "Molecular insights into the regulation of coenzyme selection in NADP-specific glutamate dehydrogenase from *Saccharomyces cerevisiae*"

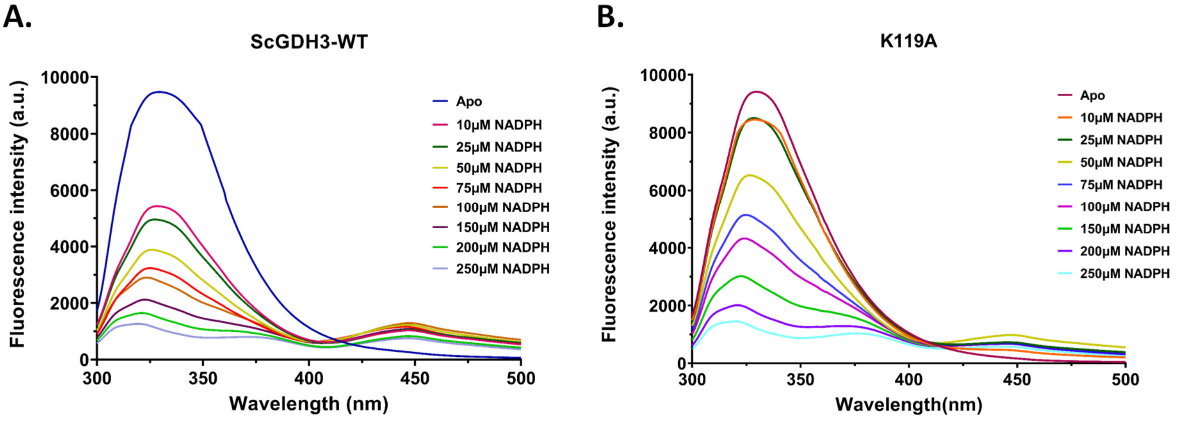


**Figure S1: Tryptophan fluorescence quenching assays of ScGDH3-WT and K119A mutant. A.** NADPH binding in ScGDH3-WT. **B.** NADPH binding in K119A mutant. Fixed concentration (6 μM) of proteins were used.

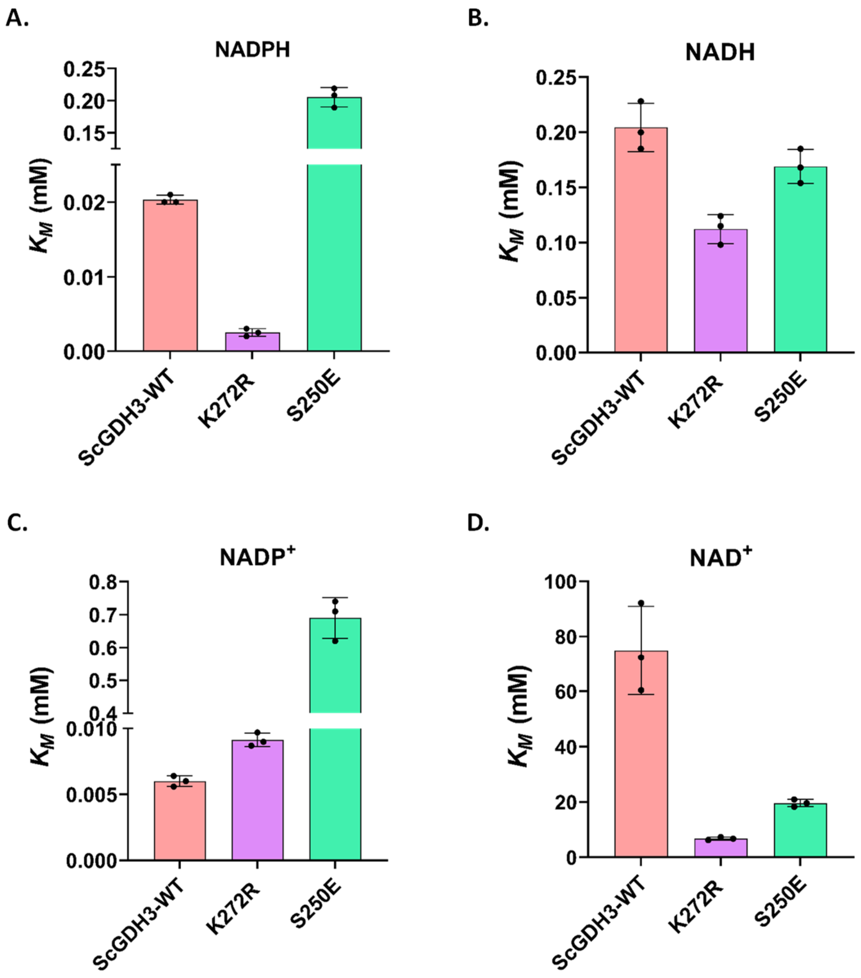


**Figure S2: Bar graphs showing $K_M$ values of ScGDH3-WT and the mutants.** $K_M$ values determined from the coenzyme saturation kinetics of ScGDH3-WT, K272R mutant and S250E mutant when **A.** NADPH is considered as coenzyme, **B.** NADH is considered as coenzyme, **C.** $NADP^+$ is considered as coenzyme, and **D.** $NAD^+$ is considered as coenzyme.

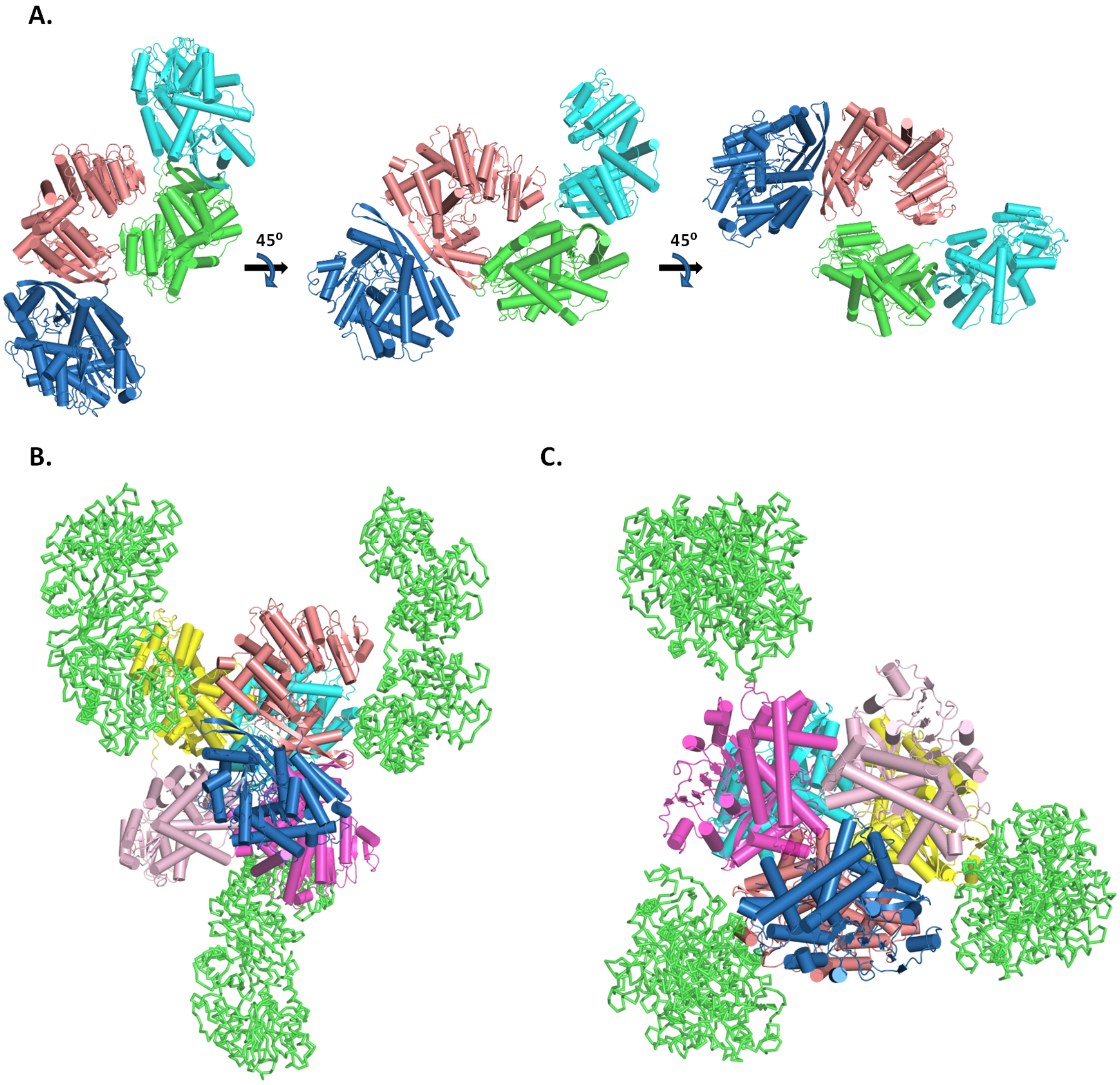


**Figure S3: Crystallographic oligomeric assembly and creation of biological hexameric assembly. A.** The crystallographic assembly shows presence of four subunits in the asymmetric unit. Different orientations of tetrameric (crystallographic) assembly are shown where subunits are represented in cartoon style (with cylindrical helices). **B.** Side view of the biological (hexameric) assembly. **C.** Top view of the biological (hexameric) assembly. Six subunits in the hexameric assembly are represented in cartoon style (with cylindrical helices) and the rest of the subunits in the crystal are colored green and represented in ribbon style.

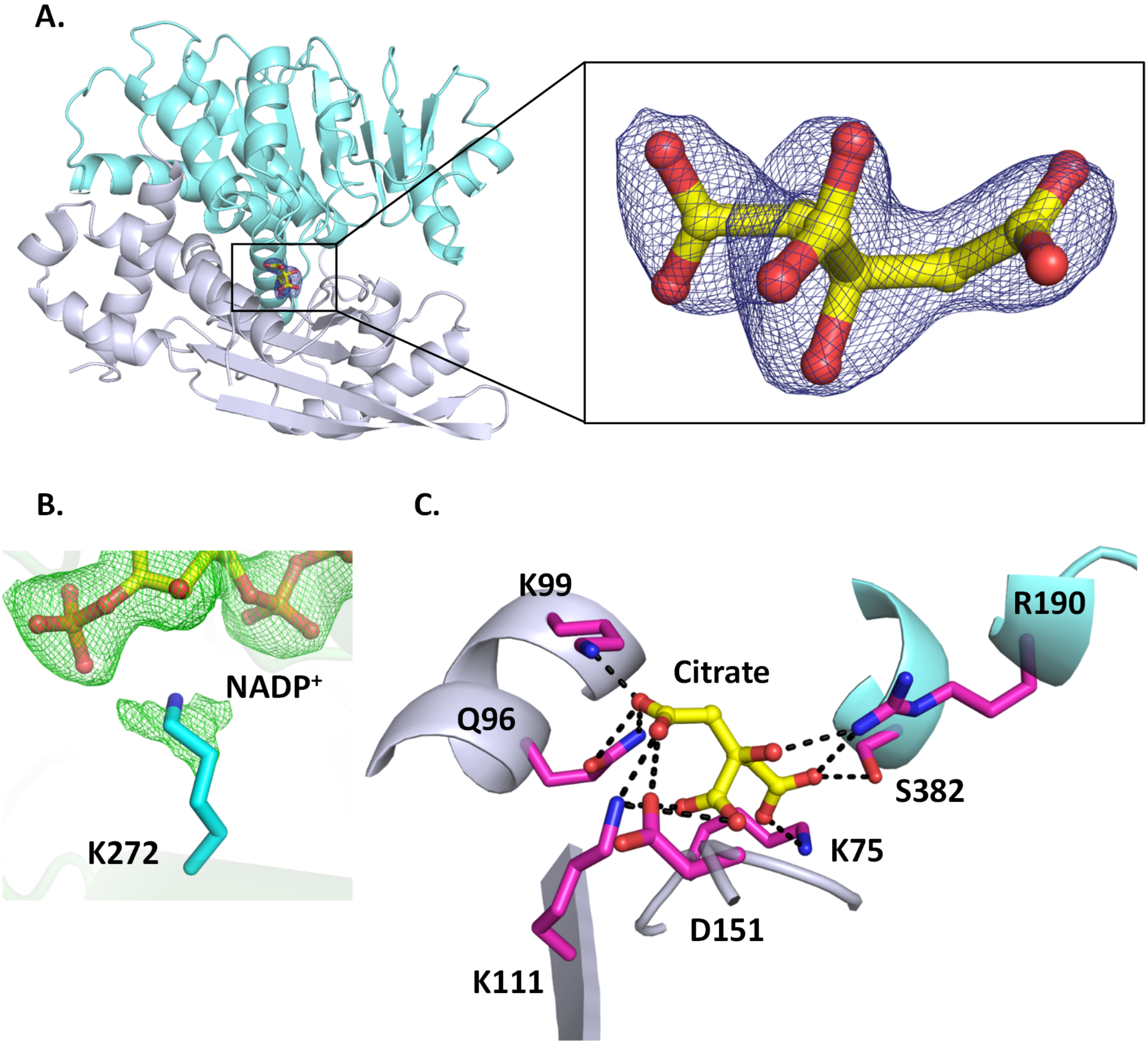


**Figure S4: Electron densities of citrate binding and mutation in ScGDH3-K272R structure. A.** Citrate (represented in ball and sticks, yellow) is present in the substrate binding site, located at the core of the cleft between domain I and domain II. Zoomed in view of the $2F_o$-$F_c$ map as light blue mesh and contoured at 1.0 σ level is shown around the bound citrate. **B.** The positive $F_o$-$F_c$ omit map (green mesh, contoured at 3.0 σ) around K272 confirms K272R mutation. **C.** The residues interacting with citrate are shown in sticks (magenta). Most of the residues are from domain I (secondary structure in cartoon, blue white). R190 and S382 belong to domain II (secondary structure in cartoon, aquamarine).

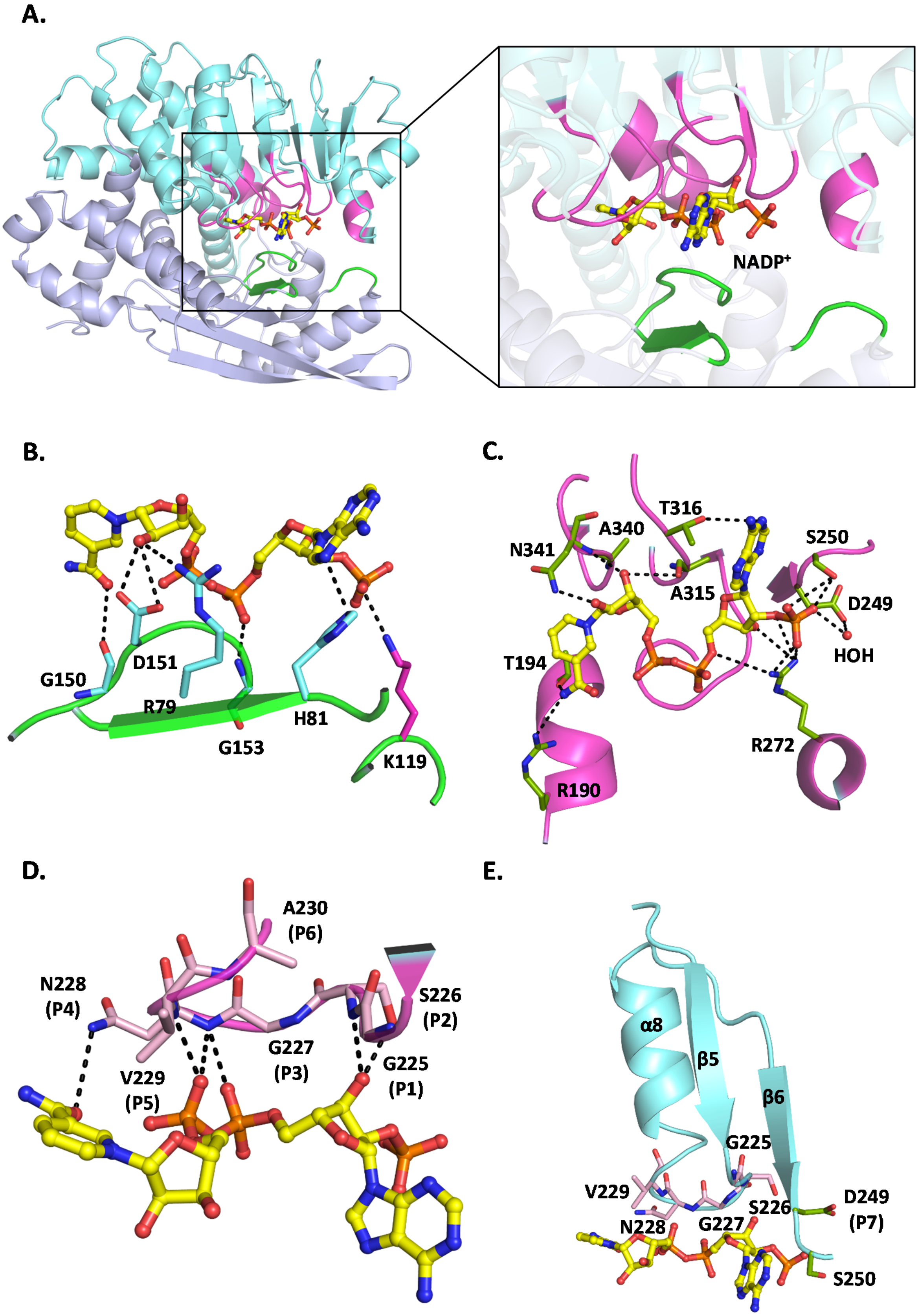
A.
NADP+
B.
G150
D151
R79
G153
H81
K119
C.
T316
A340
N341
S250
A315
D249
HOH
T194
R272
R190
D.
A230
(P6)
N228
(P4)
S226
(P2)
G227
(P3)
V229
(P5)
G225
(P1)
E.
α8
β5
β6
G225
V229
S226
D249
(P7)
N228
G227
S250

**Figure S5: Stabilization of $NADP^+$ in the coenzyme binding pocket of ScGDH3-K272R.** **A.** Coenzyme bound subunit shown in cartoon consists of domain I (blue white) and domain II (aquamarine). Zoomed in view showing the bound $NADP^+$ (ball and sticks, yellow carbon), is stabilized by the residues from both the domains. The secondary structural elements that provide the interacting residues to stabilize the coenzyme, are colored green for domain I and light magenta for domain II. **B.** The residues from domain I interacting with $NADP^+$ are represented in sticks (cyan). K119 (magenta) is the only domain I residue that directly interacts with 2´$PO_4$ moiety of $NADP^+$. **C.** The residues from domain II interacting with $NADP^+$ are represented in sticks (splitpea). Only D249 is showing a water mediated interaction with 2´$PO_4$ moiety. **D.** G225-A230 (P1-P6) residues (light pink) are part of consensus sequence "GXGXXG/A". These residues help in stabilization of the coenzyme inside the binding pocket. **E.** The modified Rossmann fold (β-α-β motif) shown in cartoon (cyan). The motif contains the consensus sequence and extremely conserved negatively charged residue (D249, P7). The side chain of D249 is pointed away from the 2´$PO_4$ moiety, as reported for other NADP-specific GDHs (Son *et al.*, 2015; Prakash *et al.*, 2018; Li *et al.*, 2021)

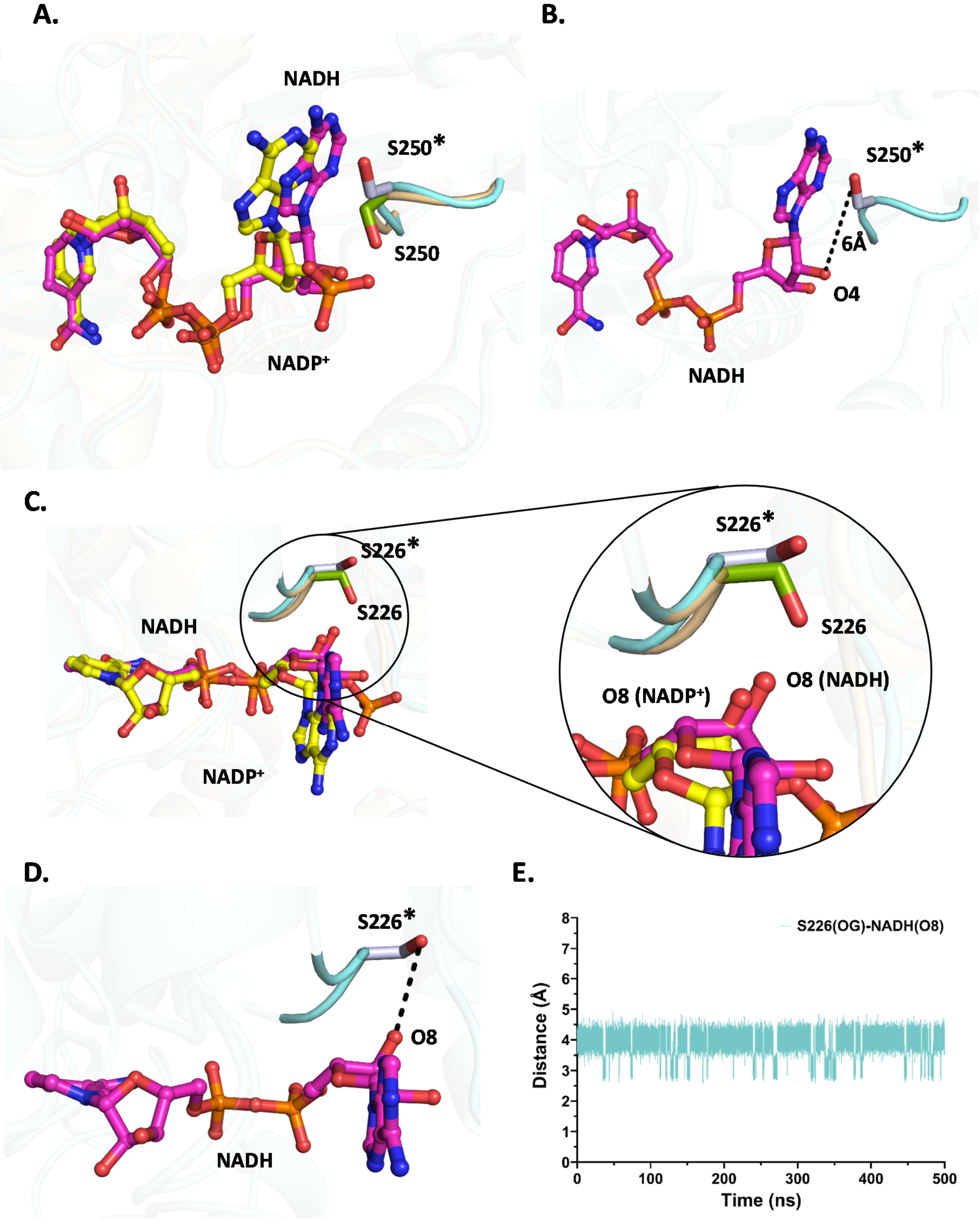
A.
NADH
S250*
S250
NADP+
B.
S250*
6Å
O4
NADH
C.
S226*
S226
NADH
NADP+
S226*
S226
O8 (NADP+)
O8 (NADH)
D.
S226*
O8
NADH
E.
S226(OG)-NADH(O8)
Distance (Å)
Time (ns)
0
1
2
3
4
5
6
7
8
0
100
200
300
400
500

**Figure S6: Alteration of the side chain orientations aids in stabilization of NADH in the coenzyme binding pocket as observed from MD simulations. A.** A significant change (~ 180$^{o}$ rotation) in the side chain orientation of S250. The side chain of S250 points towards $NADP^+$ but facing completely away when NADH is present. **B.** S250* is no longer in interacting distance (~ 6 Å) with NADH because of the alteration in side chain orientation. **C.** Slight shift in the side chain orientation between S226 (smudge) and S226* (blue white). Zoomed in view showed S226* slightly moves away from the 3′OH (O8) of adenine ribose moiety in NADH. **D**. Interaction between S226* and 3′OH (O8) group of NADH in the simulated structure. **E.** The plot is showing the trajectory (throughout 500 ns simulation) for interacting distances in between terminal –OH group (OG) of S226* and the O8 atom of NADH.

Residues of the simulated structure of the ScGDH3-K272R-NADH complex are marked by "*". The residues from the ScGDH3-K272R-$NADP^+$ crystal structure are not marked.

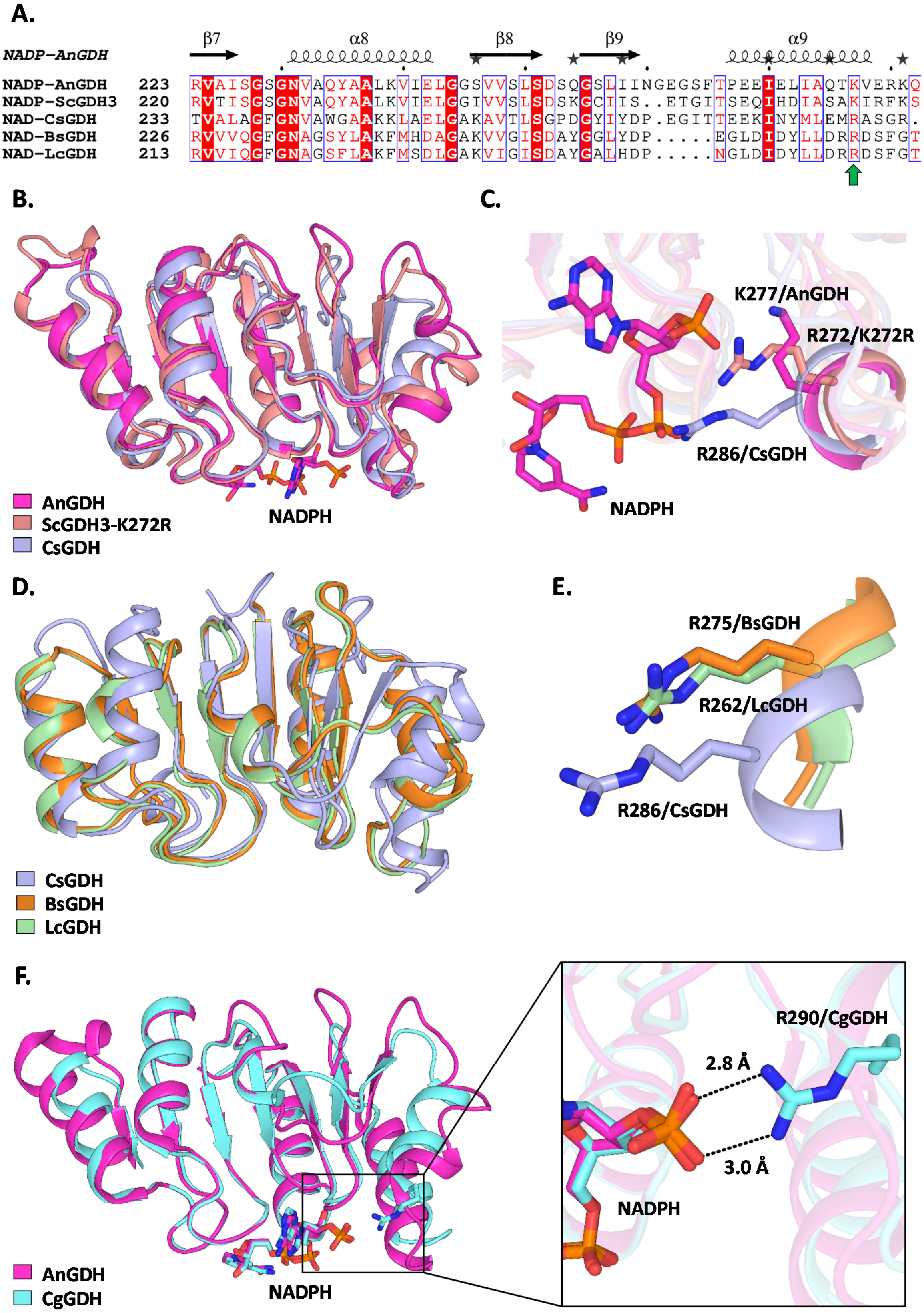
A.
β7
α8
β8
β9
α9
NADP-AnGDH
NADP-AnGDH 223 RVAISGSGNVAQYAALKVIELGGSVVSLSDSQGSLIINGEGSFTPEEIELIAQTKVERKQ
NADP-ScGDH3 220 RVTISGSGNVAQYAALKVIELGGIVVSLSDSKGCIIS..ETGITSEQIHDIASAKIRFKS
NAD-CsGDH 233 TVALAGFGNVAWGAAKKLAELGAKAVTLSGPDGYIYDP.EGITTEEKINYMLEMRASGR.
NAD-BsGDH 226 RVVVQGFGNAGSYLAKFMHDAGAKVVGISDAYGGLYDP.....EGLDIDYLLDRRDSFGT
NAD-LcGDH 213 RVVIQGFGNAGSFLAKFMSDLGAKVIGISDAYGALHDP.....NGLDIDYLLDRRDSFGT
B.
AnGDH
ScGDH3-K272R
CsGDH
NADPH
C.
K277/AnGDH
R272/K272R
R286/CsGDH
NADPH
D.
CsGDH
BsGDH
LcGDH
E.
R275/BsGDH
R262/LcGDH
R286/CsGDH
F.
AnGDH
CgGDH
NADPH
R290/CgGDH
2.8 Å
3.0 Å
NADPH

**Figure S7: Structural and sequence alignments of GDHs show evolutionary conservation of naturally occurring arginine in NAD-specific GDHs. A.** Selected portion of the sequence alignment using both NADP-specific and NAD-specific GDHs showing presence of arginine (R) in NAD-GDHs at the homologous position of lysine (K), present in NADP-GDHs (indicated by green upward arrow). The sequence alignment was performed using GDHs from *Aspergillus niger* (AnGDH), *Saccharomyces cerevisiae* (ScGDH3), *Clostridium symbiosum* (CsGDH), *Bacillus subtilis* (BsGDH) and *Lysinibacillus composti* (LcGDH). The secondary structural elements of AnGDH structure (PDB ID: 5XVX) are shown on top of the sequence alignment. **B.** Structural alignment of modified Rossmann fold and surrounding regions form domain II, contributing in coenzyme binding. The alignment is performed using AnGDH (shown in cartoon, magenta), ScGDH3-K272R (shown in cartoon, salmon) mutant and CsGDH (shown in cartoon, light blue). NADPH (represented in sticks, magenta) is present in the coenzyme binding pocket of AnGDH. **C.** Presence of K277 in AnGDH, R272 in ScGDH3-K272R and R286 in CsGDH at the homologous position. R272 and R286 protrude more inside the coenzyme binding pocket and reduce the pocket volume compared to K277, showing how arginine can stabilize NAD(H) in CsGDH and K272R mutant. **D.** Structural alignment of modified Rossmann fold and surrounding regions form domain II, among NAD-specific CsGDH (shown in cartoon, light blue), BsGDH (shown in cartoon, bright orange) and LcGDH (shown in cartoon, pale green). **E.** Structural evidence of the presence of conserved arginine (R286 in CsGDH, R275 in BsGDH and R262 in LcGDH; represented in sticks) in these three NAD-GDHs. **F.** Structural alignment of modified Rossmann fold and surrounding regions form domain II, in between NADP-specific AnGDH (shown in cartoon, magenta) and NADP-specific *Corynebacterium glutamicum* GDH (CgGDH) (shown in cartoon, aquamarine). Both have coenzymes present inside the binding pocket. Zoomed in view shows direct interaction between 2´$PO_4$ moiety and R290 (represented in sticks) from CgGDH, suggesting arginine can also stabilize NADP(H) inside the coenzyme binding pocket.

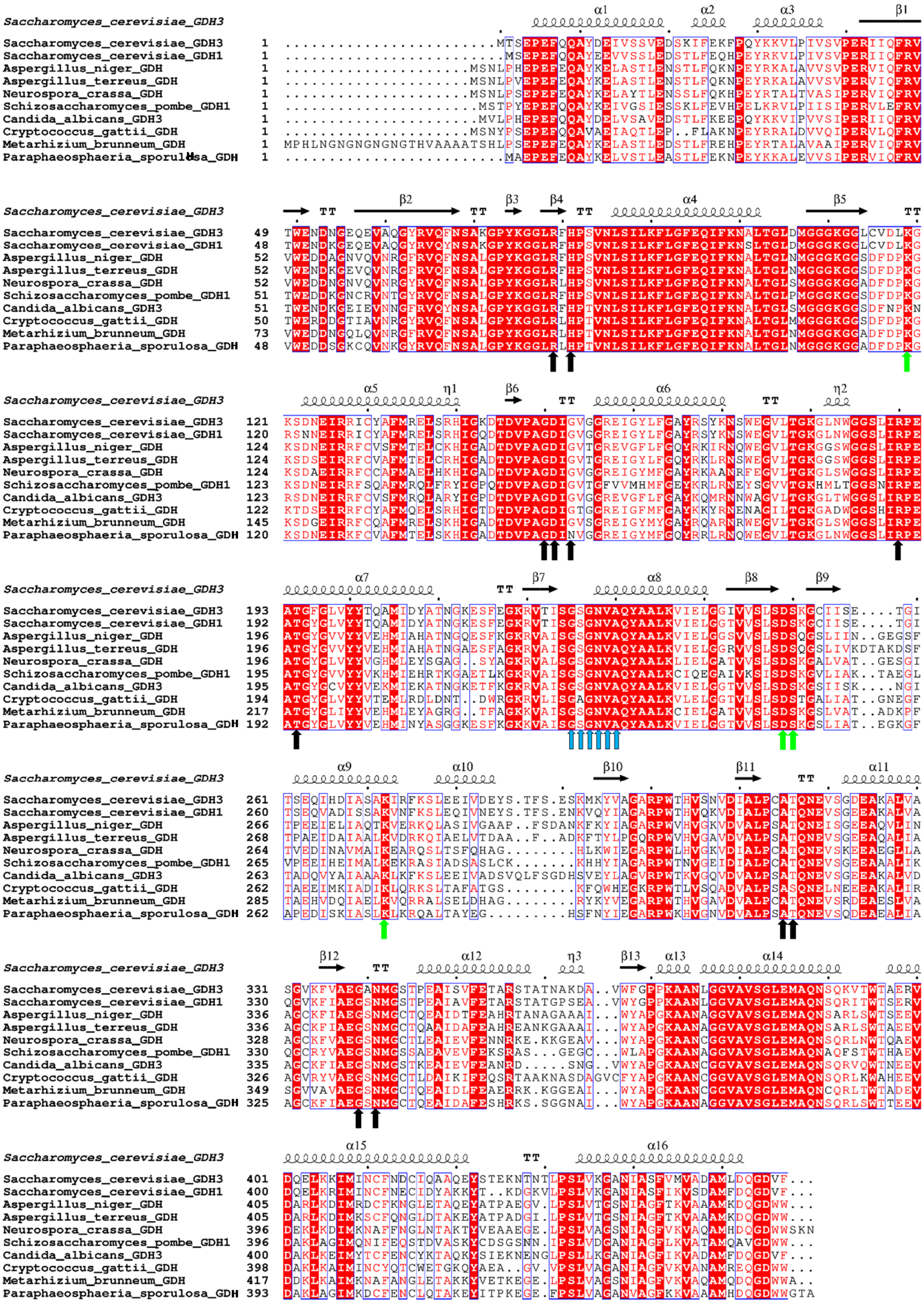

**Figure S8: Sequence alignment of NADP-specific GDHs from different fungal species.** The upward arrows in different colors (black, green, light blue), indicate residues from domain I and domain II which help in stabilization of the coenzyme inside coenzyme binding pocket. All these residues are extremely conserved. The residues indicated by yellow upward arrows are part of the consensus sequence "GXGXXG/A" (P1-P6), present in all dehydrogenase enzymes including GDHs. Four residues (K119, D249, S250 and K272) indicated by green upward arrows are involved in direct/indirect interaction with 2´PO4 moiety of NADPH. D249 is highly conserved acidic residue among dehydrogenases and is denoted as "P7". Residues are numbered with respect to *Saccharomyces cerevisiae* GDH3 (ScGDH3). The secondary structural elements of ScGDH3-K272R structure (PDB ID: 47UL) are shown on top of the sequence alignment.

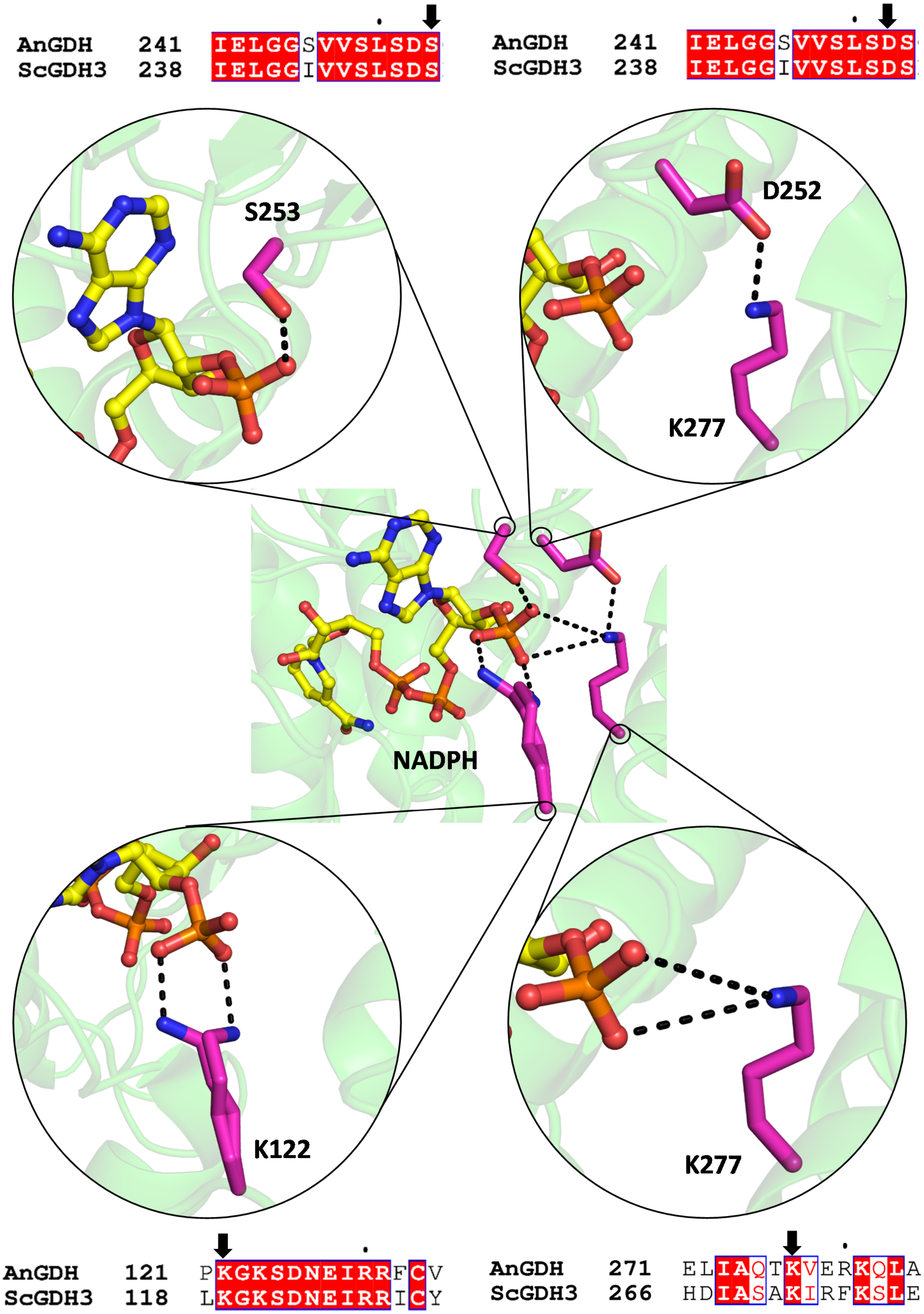
AnGDH 241 IELGGSVVSLSDS
ScGDH3 238 IELGGIVVSLSDS
AnGDH 241 IELGGSVVSLSDS
ScGDH3 238 IELGGIVVSLSDS
S253
D252
K277
NADPH
K122
K277
AnGDH 121 PKGKSDNEIRRFCV
ScGDH3 118 LKGKSDNEIRRICY
AnGDH 271 ELIAQTKVERKQLA
ScGDH3 266 HDIASAKIRFKSLE

**Figure S9: Molecular basis of binding of the 2´$PO_4$ moiety of NADPH in *Aspergillus niger* GDH (AnGDH) and development of the hypothesis of possible mutation in ScGDH3 to alter its coenzyme specificity.** Zoomed in views showing direct interaction with S253 (top-left), K122 (bottom-left), and K277 (bottom-right). D252 is pointed away from the 2´$PO_4$ moiety and is stabilized by K277 (top-right). K122 is the only residue from domain I which interacts with the 2´$PO_4$ moiety (Prakash *et al.*, 2018). Sequence alignment results show all these residues are conserved in NADP(H)-specific ScGDH3.

**Table S1: Primers for mutagenesis studies on ScGDH3**

| Primer | Sequence (5´ to 3´) |
|---|---|
| ScGDH3-K272R_forward | TCCGCC**<u>CGG</u>**ATCCGTTTCAAGTCGTTAGAGG |
| ScGDH3-K272R_reverse | GGAT**<u>CCG</u>**GGCGGAAGCGATATCGTGAATTTGC |
| ScGDH3-K272A_forward | TCCGCC**<u>GCT</u>**ATCCGTTTCAAGTCGTTAGAGG |
| ScGDH3-K272A_reverse | GGAT**<u>AGC</u>**GGCGGAAGCGATATCGTGAATTTGC |
| ScGDH3-S250E_forward | TTATCCGAT**<u>GAG</u>**AAGGGGTGCATCATCTCTGAG |
| ScGDH3-S250E_reverse | GCACCCCTT**<u>CTC</u>**ATCGGATAAAGACACCACAATACC |
| ScGDH3-K119A_forward | GACTTG**<u>GCT</u>**GGCAAGTCTGACAACGAGATCAGAAGG |
| ScGDH3-K119A_reverse | CTTGCC**<u>AGC</u>**CAAGTCCACACACAGGCCACCCTTACC |
| ScGDH3-K119F_forward | GACTTG**<u>TTC</u>**GGCAAGTCTGACAACGAGATCAGAAGG |
| ScGDH3-K119F_reverse | CTTGCC**<u>GAA</u>**CAAGTCCACACACAGGCCACC |

**Media S1:** Formation of crystallographic biological (hexameric) assembly from adjacent asymmetric units containing tetramer. The hexameric assembly, in the center, is shown in cartoon (with cylindrical helices) and each subunit is colored differently. Initially, the rest of the molecules from adjacent asymmetric units are shown in ribbon (green). Two molecules from each asymmetric unit contribute to the formation of biological assembly. Remaining two molecules from each asymmetric unit, are shown in cartoon (with cylindrical helices, smudge). This media has been generated in PyMol.
